# Human MAIT cells show clonal diversity but transcriptional and functional homogeneity

**DOI:** 10.1101/2022.02.26.482031

**Authors:** Lucy C. Garner, Ali Amini, Michael E.B. FitzPatrick, Nicholas M. Provine, Paul Klenerman

## Abstract

Mucosal-associated invariant T (MAIT) cells are considered to have limited clonal diversity. In contrast, recent studies suggest the presence of functionally distinct subsets. We investigated this model through single-cell analysis of the MAIT cell TCR repertoire and transcriptional profile in human blood and liver. Further, we developed functional RNA-sequencing (fRNA-seq), an approach to integrate cellular function and TCR clonotype at a single-cell level following differential stimulation. MAIT cells showed surprising clonal diversity, with TCR repertoires shared across tissues but unique to individuals. Functional diversity within resting MAIT cells was low and largely related to tissue site. MAIT cells displayed distinct transcriptional responses to in vitro TCR and cytokine stimulation, with cells positioned along gradients of activation. Clonal origin influenced both resting and activated transcriptional profiles. Overall, MAIT cells exhibit diverse donor-specific TCR repertoires which, along with tissue and activation context, influence their phenotype and function.

## Introduction

Mucosal-associated invariant T (MAIT) cells are a population of innate-like T cells, abundant in human blood (1-10% of T cells) and peripheral tissues, particularly the liver (2-50% of T cells) and mucosal sites^1^. Unlike conventional T cells that express diverse MHC-restricted peptide-specific T cell receptors (TCR), MAIT cells express a semi-invariant Vα7.2-Jα33/12/20 (*TRAV1-2-TRAJ33*/*12*/*20*) TCR specific for bacterial- and yeast-derived riboflavin metabolites presented by the MHC Class I-like molecule MR1^1^. In addition to TCR-dependent activation, MAIT cells can be activated independent of their TCR by cytokines such as IL-12 and IL-18^2^. Upon activation, they secrete type 1 and/or type 17 cytokines, and exhibit cytotoxic activity^3, 4^.

A major outstanding question in the MAIT cell field is whether human MAIT cells comprise transcriptionally and functionally distinct subsets. Alterations in MAIT cell frequency, phenotype and function occur in a broad range of human infectious and inflammatory diseases, and mouse models indicate both protective and pathogenic roles^5^. Improved knowledge of the characteristics and structure of the MAIT cell population in health could aid the development of therapeutics targeting specific subsets or functions during disease.

In human blood, MAIT cells appear relatively homogeneous, exhibiting a predominantly CD8^+^ effector-memory phenotype, and characteristic expression of surface molecules (e.g. CD161, CD26) and transcription factors (e.g. T-bet, RORγt, PLZF)^1^. However, some variability in phenotype and function has been identified, including differences between CD8^+^, CD4^+^ and double-negative (DN, CD4^-^CD8^-^) cells^6–8^, and variable surface expression of innate immune receptors^8, 9^. Despite universal expression of the type 17 transcription factor RORγt, only a small percentage (< 5%) of human MAIT cells produce IL-17 ex vivo^3^. This may be explained by the presence of a distinct type 17 subset. Consistent with this hypothesis, MAIT1 and MAIT17 subsets are present in mice^10–12^, and related invariant natural killer T (iNKT) cells comprise multiple subsets^13, 14^. However, the existence of analogous transcriptionally distinct MAIT cell subsets has not been proven in humans.

MAIT cell function is altered by tissue localisation and stimulation. Compared with blood, gut and liver MAIT cells display a more activated and tissue-resident transcriptome^12, 15–17^, while female genital tract^18^ and oral mucosal^19^ MAIT cells are skewed towards type 17 functions. In adipose tissue around 15% of MAIT cells secrete IL-10^20^, a cytokine not produced in other tissues. MAIT cells exhibit distinct transcriptional responses to TCR and cytokine stimulation^21, 22^, and produce type 2 cytokines^23^ and increased IL-17^24^ following prolonged stimulation. Whether functional differences across tissues and upon stimulation reflect the presence of multiple subsets or environment-driven plasticity in the entire population, remains unknown.

Additionally, there are questions surrounding the MAIT cell TCR repertoire, including the variability across tissues and donors, and the relationship between TCR usage and function. Some studies demonstrate similar TCR repertoires across tissues^25^, whereas others indicate differences in *TRAJ*^1^^9, 26^ or *TRBV*^27^ usage. Diverse chain usage within and between tissues could have functional implications. MAIT cells responding to stimulation with different pathogens in vitro show distinct TCR repertoires^28^. During human *Salmonella* infection, the distribution of MAIT cell clonotypes changes, and cells transduced with TCRβ chains from expanded clonotypes show a greater response to in vitro TCR stimulation compared with cells transduced with TCRβ chains from contracted clonotypes^29^. This is consistent with studies showing differential activation potential of MAIT cell clones^28^, as well as MAIT cells with different Vβ segments^9^ or CDR3β sequences^30^. Thus, several studies suggest a relationship between TCR architecture and MAIT cell function, but this has not been studied at the single-cell level with large numbers of cells.

In summary, human MAIT cells show some variation in phenotype, function and TCR repertoire. However, previous studies have examined a limited number of cells or proteins, or performed bulk RNA-seq on marker-sorted populations. Thus, it is unknown whether human MAIT cells comprise multiple functionally distinct subsets and how function relates to TCR usage. To investigate this, we performed single-cell RNA-sequencing (scRNA-seq) and single-cell TCR-sequencing (scTCR-seq) of human MAIT cells from matched blood and liver, as well as blood MAIT cells at rest and following TCR and cytokine stimulation. Our findings indicate surprising TCR diversity within human MAIT cells, in contrast with a largely conserved transcriptional program. Differences in tissue localisation, clonal origin and activation state were driving factors for the limited observed transcriptional diversity.

## Results

### MAIT cells exhibit tissue-specific transcription and regulation

To investigate MAIT cell heterogeneity using an unbiased genome-wide approach, we performed scRNA-seq and scTCR-seq of sorted MAIT cells (CD3^+^MR1/5-OP-RU^+^) from matched human blood (n = 3) and liver (n = 4) using 10x Genomics technology (Extended Data Fig. 1). Conventional memory T (T_mem_; CD3^+^MR1/5-OP-RU^-^CCR7^-^) cells were also analysed in two donors.

In total, 20,417 MAIT cells were analysed post-filtering. MAIT cells comprised 11 clusters (Fig. 1a), with blood and liver cells occupying different regions of the uniform manifold approximation and projection (UMAP) (Fig. 1b). Individual clusters mostly contained cells from a single tissue (Fig. 1c) but multiple donors (Extended Data Fig. 2a). Blood and liver MAIT cells differentially expressed 174 genes (Supplementary Table 1; Extended Data Fig. 2b), the majority of which were upregulated in the liver (151 genes). Liver-enriched genes encoded tissue-residency markers (e.g. *CD69*), TCR-induced transcription factors (e.g. *NR4A1*, *EGR1*), effector cytokines (e.g. *IFNG*, *TNF*), chemokines/chemokine receptors (e.g. *CCL3*, *CXCR6*) and heat-shock proteins (e.g. *HSPA1A*/*B*). Some genes such as *CD69* and *CXCR6* showed uniformly higher expression in liver compared with blood, while others such as *CCL3* and *CCL20* were enriched in specific clusters (Fig. 1d).

**Fig. 1.**
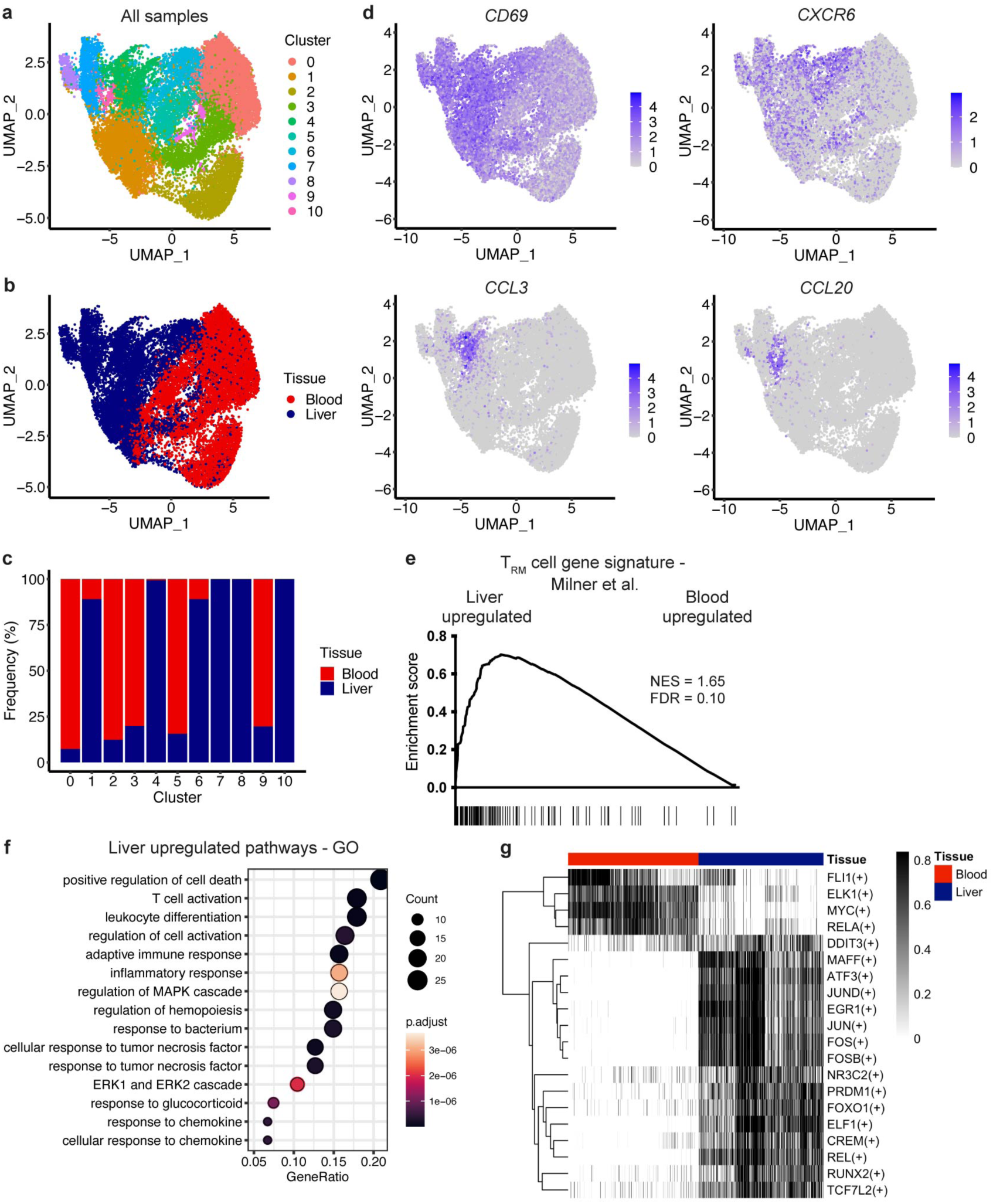
Liver MAIT cells exhibit an activated, tissue-resident transcriptional and regulatory profile. **a**, UMAP of blood and liver MAIT cells coloured by the 11 identified clusters. **b**, UMAP coloured by tissue. **c**, Proportion of cells in each cluster from the blood and liver. **d**, UMAPs coloured by the expression of *CD69*, *CXCR6*, *CCL3* and *CCL20*. **e**, Gene set enrichment analysis (GSEA) of liver compared with blood MAIT cells using a tissue-resident memory T (T_RM_) cell gene signature from Milner and colleagues^31^. NES = normalised enrichment score, FDR = false discovery rate. **f**, Pathway analysis on the genes significantly upregulated in liver compared with blood MAIT cells. Top 15 non-redundant gene ontology (GO) terms are shown. **g**, Heatmap showing the activity (row-scaled AUCell scores) of the top 20 (based on fold change) significantly differentially active regulons between blood and liver.

To establish whether liver MAIT cells showed a more tissue-resident transcriptional profile compared with blood MAIT cells, we performed Gene Set Enrichment Analysis (GSEA) using a mouse tissue-resident memory T (T_RM_) cell gene signature published by Milner and colleagues^31^. The T_RM_ cell signature was highly enriched in the liver (Fig. 1e). The 70 leading edge genes included *CD69*, *TNF*, *HAVCR2* (TIM-3), *ICOS* and *FOS*. Enrichment of tissue-residency genes was confirmed using a second T_RM_ cell signature^32^ (Extended Data Fig. 2c). Other liver-enriched pathways were associated with inflammation, and T cell activation and differentiation (Fig. 1f).

To examine the regulation of tissue-specific gene expression, we used pySCENIC^33, 34^ to discover transcription factor regulons – modules of genes predicted to be regulated by a given transcription factor. We identified 134 high confidence regulons, 122 of which were differentially active in blood and liver (Supplementary Table 2). Most of the top 20 differentially active regulons showed increased activity in the liver (Fig. 1g). These included PRDM1 (encodes Blimp-1), the TCR-induced transcription factor EGR1, REL (NF-kB family) and multiple members of the AP-1 family (FOS, FOSB, JUN, JUND, ATF3, MAFF) (Extended Data Fig. 2d). Genes regulated by REL and several AP-1 factors, particularly JUND and ATF3, were significantly enriched for cell activation and adhesion pathways (Extended Data Fig. 2e-g).

### MAIT cells show diverse TCRβ chain usage, despite a restricted TCRa chain repertoire

Having demonstrated that blood and liver MAIT cells were transcriptionally distinct, we next investigated whether the TCR repertoire was tissue specific. Previous analyses were limited by scale (e.g. examination of T cell clones or single-cell sequencing of small cell numbers) or depth (e.g. flow cytometry-based analyses or bulk RNA-sequencing [RNA-seq])^8, 26, 35–37^. Thus, our dataset of > 20,000 MAIT cells from multiple donors and matched tissues provided a unique opportunity to examine TCR repertoire characteristics and diversity across tissues and individuals.

TCR clonotypes were defined as cells with identical TCR gene segment usage, and CDR3α and CDR3β nucleotide sequences. MAIT cell clonotypes were required to have a *TRAV1-2* TCRα chain. MAIT cells were highly oligoclonal (Fig. 2a), with oligoclonality comparable across donors and tissues. Surprisingly, the degree of MAIT cell oligoclonality was analogous to that of T_mem_ cells (Fig. 2b). MAIT cell clonotypes defined using only the TCRα chain (hereafter TCRα only clonotypes) were more oligoclonal than those defined using only the TCRβ chain (hereafter TCRβ only clonotypes) (Fig. 2c; Extended Data Fig. 3a-c). Namely, the Gini coefficient (a measure of inequality) was significantly increased, and the Shannon diversity index was significantly decreased, for TCRα only relative to TCRβ only clonotypes. At the population level, MAIT cell TCRα chain pairing was quite promiscuous, with up to 40% of detected TCRα chains paired with more than one unique TCRβ chain – generating multiple clones with identical TCRα chains (Fig. 2d). Conversely, most TCRβ chains paired with a single TCRα chain (Fig. 2e). This contrasted with T_mem_ cells, where TCRαβ pairings were essentially unique (Extended Data Fig. 3d,e).

**Fig. 2.**
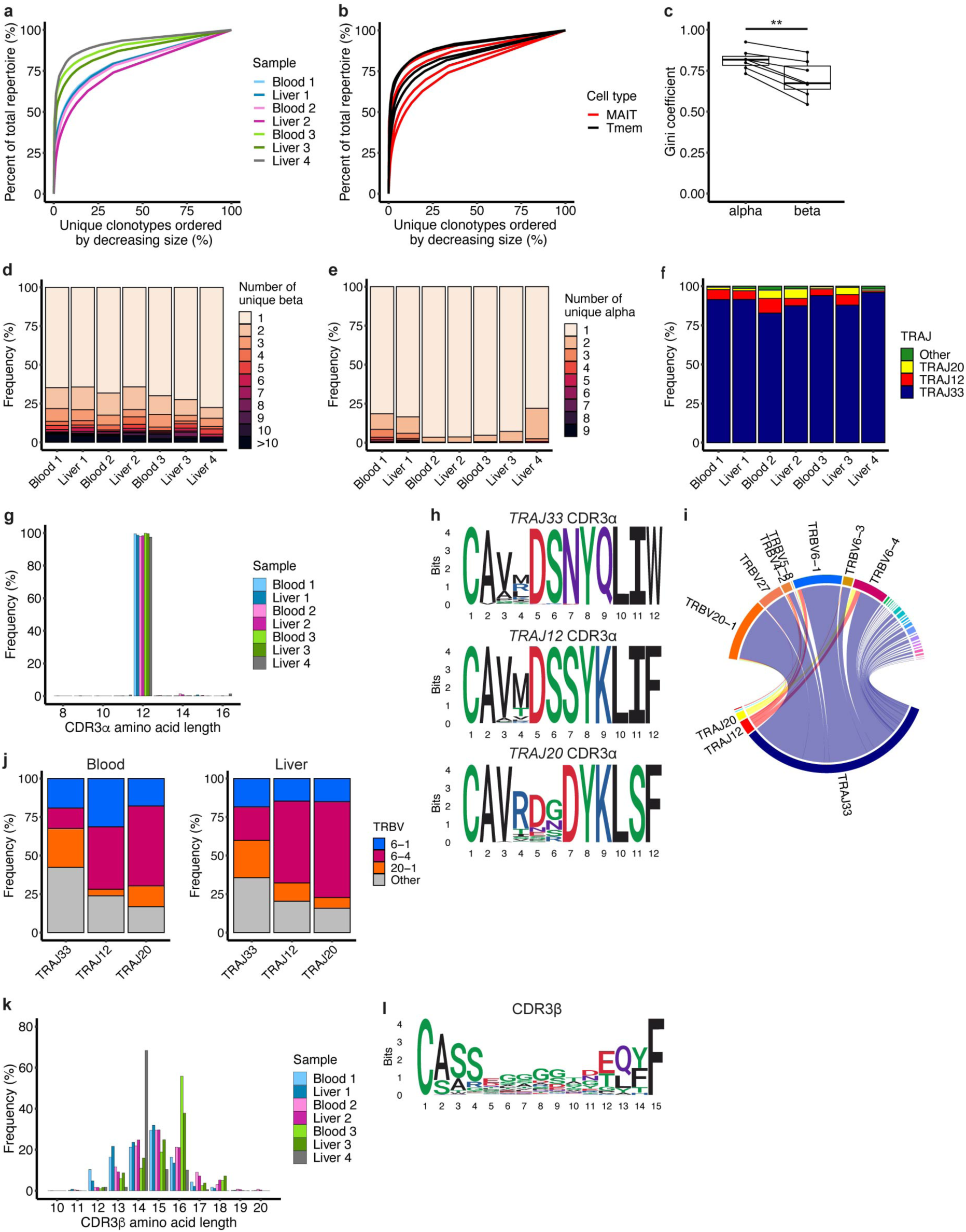
AIT cells have a restricted TCRα but diverse TCRβ repertoire. **a**, Line plot demonstrating the clonality of the MAIT cell TCRαβ repertoire coloured by sample. **b**, Line plot comparing MAIT (red) and T_mem_ (black) cell TCRαβ clonality (blood and liver samples from donors 2 and 3). **c**, Gini coefficient for MAIT cell TCRα only and TCRβ only clonotypes. Paired t-test, ** p < 0.01. **d**,**e**, MAIT cell TCR chain pairing at the population level. Number of unique TCRβ chains paired with a single TCRα chain (**d**), or TCRα chains paired with a single TCRβ chain (**e**), in MAIT cells from each sample. **f**, Proportion of MAIT cells expressing *TRAJ33*, *TRAJ12*, *TRAJ20* and other *TRAJ* gene segments in each sample. **g**, MAIT cell CDR3α amino acid length in each sample. **h,** Sequence logos generated from all MAIT cell *TRAJ33*, *TRAJ12* or *TRAJ20* CDR3α amino acid sequences of length 12. **i**, Circos plot showing the average use of *TRAJ* and *TRBV* gene segments amongst MAIT cells combined from all seven samples. **j**, Proportion of *TRAJ33*, *TRAJ12* and *TRAJ20* MAIT cell TCRs with *TRBV6-1*, *TRBV6-4*, *TRBV20-1* and other *TRBV* gene segments. **k**, MAIT cell CDR3β amino acid length in each sample. **l**, Sequence logo generated from all MAIT cell CDR3β amino acid sequences of length 15.

*TRAJ33*, *TRAJ12* and *TRAJ20* were used by 90%, 5% and 3% of TCRs, respectively (Fig. 2f), with no significant difference between blood and liver (Extended Data Fig. 3f). Other *TRAJ* gene segments (27 in total) were each present at a frequency of less than 0.5%. CDR3α sequences were 12 amino acids long (Fig. 2g), conserved across donors and tissues, and included the tyrosine 95 residue (CDR3α position eight) required for MAIT cell activation^30, 38, 39^ (Extended Data Fig. 3g). Use of different *TRAJ* gene segments resulted in systematic differences in CDR3α sequence, and the number of N and P nucleotide substitutions (Fig. 2h; Extended Data Fig. 3h,i).

MAIT cells preferentially expressed *TRBV6-1*, *TRBV6-4,* and *TRBV20-1* TCRβ gene segments^26, 35, 40^ (Fig. 2i), with no significant difference in *TRBV* usage between blood and liver (Extended Data Fig. 3j). However, *TRBV* usage was diverse, with 48 *TRBV* gene segments expressed. Enrichment of specific chains showed wide variation between donors, often driven by large expanded clonotypes (Extended Data Fig. 3k-n).

As demonstrated in Fig. 2h, *TRAJ* usage influenced CDR3α sequence. Studies of small numbers of clones or TCR sequences suggest *TRAJ* usage could affect TCRαβ pairing^26, 35^. Therefore, we examined *TRBV* usage by *TRAJ33*, *TRAJ12* and *TRAJ20* TCRs. *TRAJ12 and TRAJ20* TCRα chains showed increased pairing with *TRBV6-4* TCRβ chains compared with *TRAJ33*, while *TRAJ33* TCRα chains showed increased pairing with *TRBV20-1* TCRβ chains (Fig. 2j; Extended Data Fig. 3o,p).

CDR3β length and sequence were highly variable, in contrast with the CDR3α region (Fig. 2k,l; Extended Data Fig. 3q). Approximately 25% of CDR3β sequences were each of 14, 15 and 16 amino acids long. Due to large individual clonotypes, CDR3β lengths were skewed towards 14 and 16 amino acids in donors 4 and 3, respectively.

### The MAIT cell TCR repertoire is highly shared between blood and liver, but unique to each individual

Given the similarities in TCRα and TCRβ chain usage between matched tissues, we hypothesised that blood and liver MAIT cells might show a large degree of repertoire sharing. For each individual, over 70% of MAIT cells from matched blood and liver belonged to a shared TCR clonotype (Fig. 3a). For example, 71% of cells (3,554) from donor 2 belonged to a clonotype detected in blood and liver. Matched blood and liver T_mem_ cells showed similarly high clonal overlap (Fig. 3b).

**Fig. 3.**
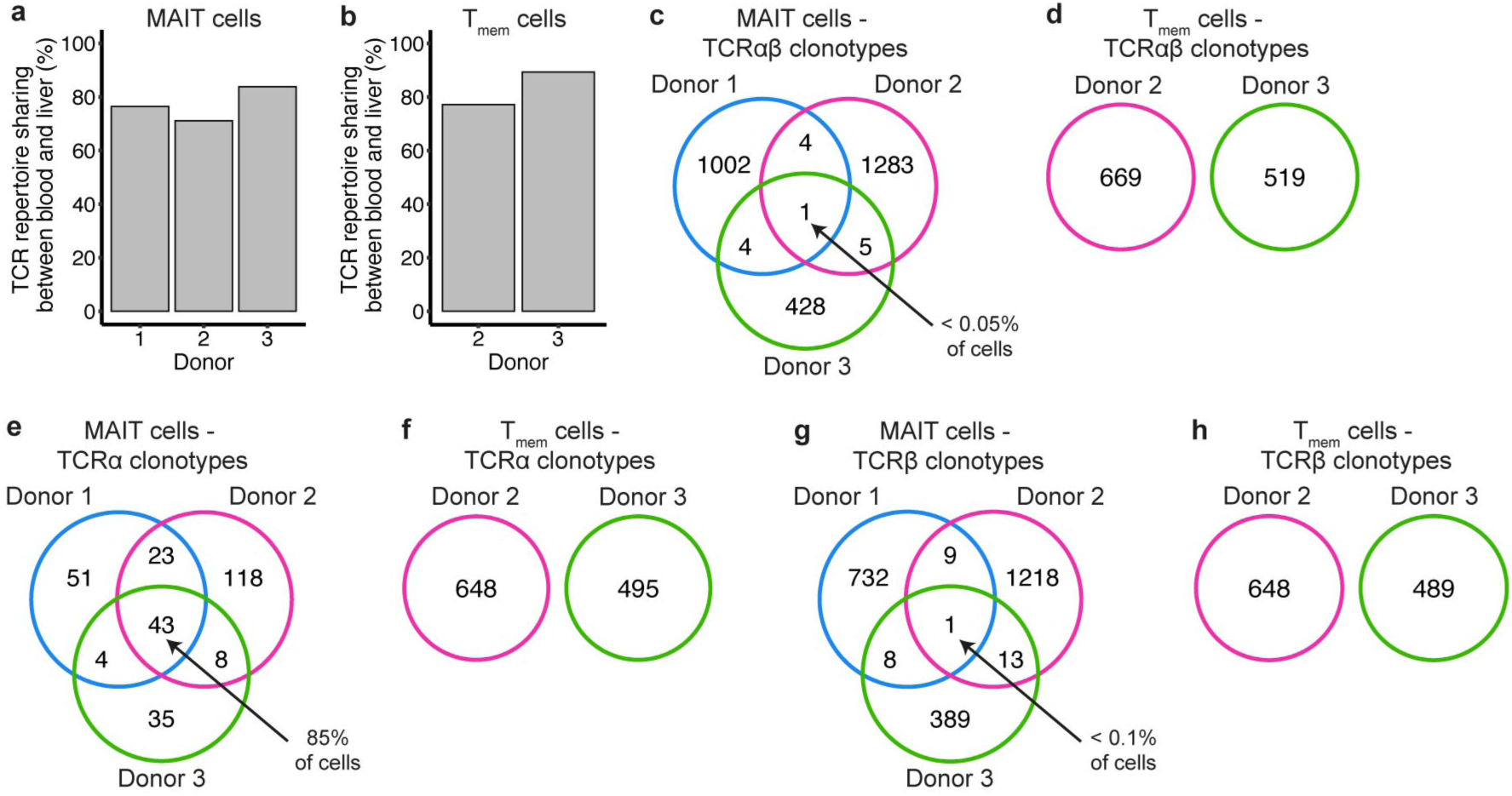
The MAIT cell TCRαβ repertoire is essentially unique to each individual but shared across tissues. **a**,**b**, Percentage of MAIT (**a**) or T_mem_ (**b**) cells belonging to a TCRαβ clonotype shared between matched blood and liver. **c**,**d**, Overlap of MAIT (**c**) or T_mem_ (**d**) cell functional TCRαβ clonotypes between donors. **e**,**f**, Overlap of MAIT (**e**) or T_mem_ (**f**) cell functional TCRα clonotypes between donors. **g**,**h**, Overlap of MAIT (**g**) or T_mem_ (**h**) cell functional TCRβ clonotypes between donors. Numbers in Venn diagrams (**c**-**h**) indicate the number of clonotypes.

We next examined repertoire sharing across donors. Clonotypes were defined based on CDR3α and CDR3β amino acid sequences as opposed to nucleotide sequences (hereafter functional clonotypes), as it is the protein structure that determines the functionality of the TCR. Despite use of a semi-invariant TCR, 98% of MAIT cells belonged to a donor-specific clonotype, with a single clonotype shared between all three donors (Fig. 3c). T_mem_ cell functional clonotypes showed no overlap between the two donors analysed (Fig. 3d).

Given the restricted TCRα chain repertoire of MAIT cells, we reasoned that functional clonotypes defined based on the TCRα chain only (hereafter functional TCRα clonotypes) may show high overlap between donors. Consistent with this hypothesis, the three donors shared 43 functional TCRα clonotypes comprising 85% (11,380/13,398) of MAIT cells (Fig. 3e). In contrast, there were no shared T_mem_ cell functional TCRα clonotypes between donors (Fig. 3f). Functional TCRβ clonotypes were largely donor-specific for MAIT and T_mem_ cells (Fig. 3g,h).

Thus, distinct from T_mem_ cells, the MAIT cell TCRα chain repertoire is public and highly shared between individuals, while the TCRβ chain is almost entirely private and governs uniqueness of individual MAIT cell TCR repertoires.

### MAIT cells show limited within-tissue transcriptional heterogeneity

Our data so far indicated distinct transcriptional and regulatory profiles of blood and liver MAIT cells despite highly overlapping TCR repertoires. We next wanted to explore the extent of within-tissue heterogeneity and whether transcriptional variation was associated with clonal origin.

Blood MAIT cells combined from all donors comprised eight clusters (Fig. 4a). Except for cluster 6 that was donor-specific, there was a roughly equal proportion of cells from the three donors in each cluster (Extended Data Fig. 4a). Transcriptional diversity between clusters was low (Fig. 4b; Supplementary Table 3a). Seven out of eight clusters had three or fewer cluster markers with a fold change (FC) > 1.5. Cluster markers did not reflect the presence of known, functionally relevant, T cell subsets. Using gene signatures from mouse MAIT cells^41, 42^, we were unable to define human MAIT1 and MAIT17 subsets (Fig. 4c). Two clusters were defined by recognisable pathways. A small cluster of MAIT cells (cluster 7; 32 cells) displayed increased expression of *STAT1*, *STAT2* and interferon-stimulated genes (Fig. 4b; Extended Data Fig. 4b). Cluster 2 showed potential functional relevance with enriched expression of granulysin and granzymes (Fig. 4d; Extended Data Fig. 4c). This cluster did not simply reflect activated MAIT cells, as *GZMB* and *GZMH* (lowly expressed in resting MAIT cells^43, 44^) were only expressed by a small percentage of cells.

**Fig. 4.**
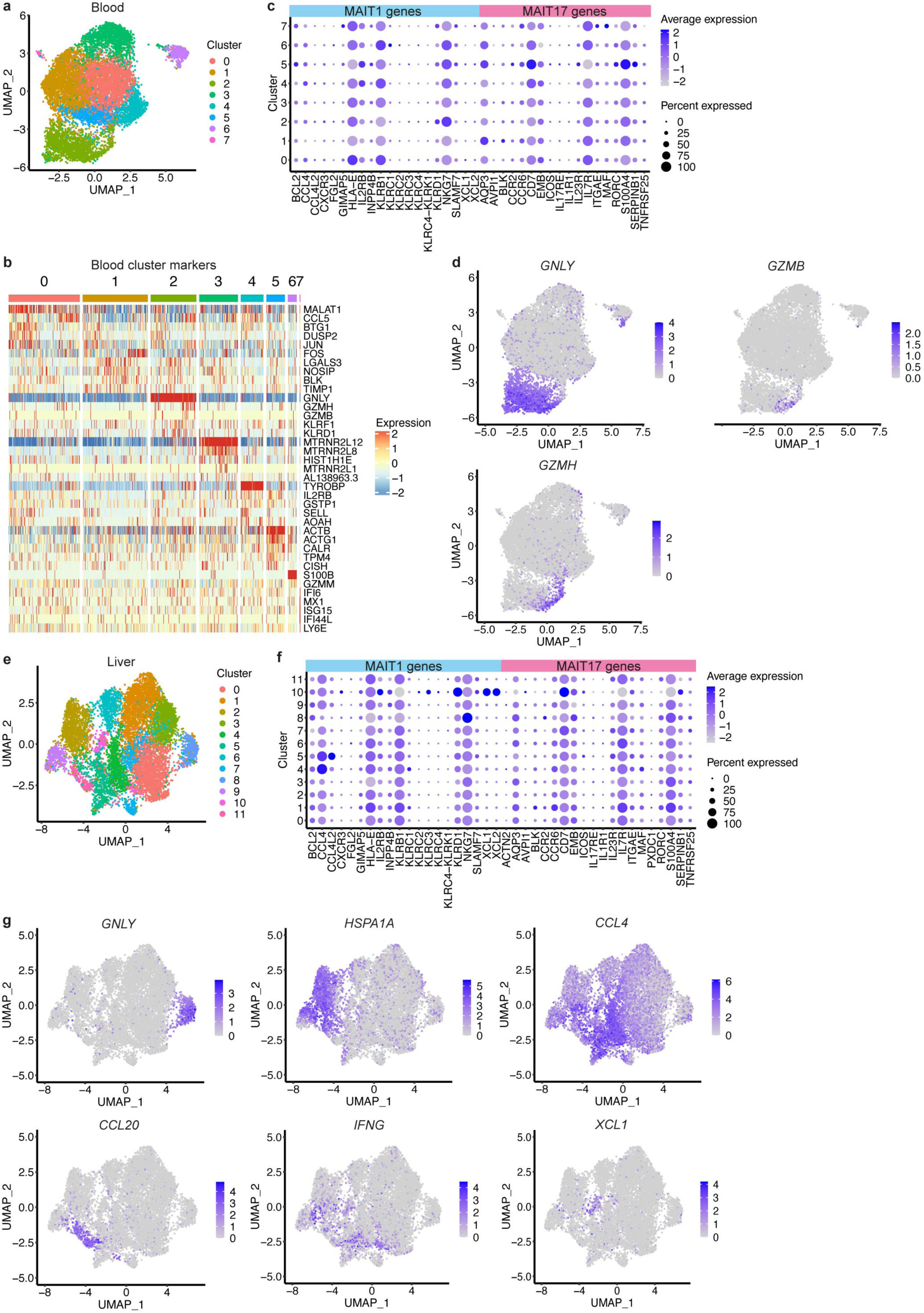
MAIT cells within the blood and liver show minimal transcriptional heterogeneity. **a**, UMAP of blood MAIT cells coloured by the eight identified clusters. **b**, Heatmap showing row-scaled log-transformed normalised expression of the top five or all (if < 5) significant marker genes for each blood MAIT cell cluster. **c**, Expression of MAIT1 and MAIT17 genes in blood MAIT cell clusters. Dot size indicates the percentage of cells expressing the gene and dot colour indicates the level of expression. **d**, UMAPs of blood MAIT cells coloured by the expression of *GNLY*, *GZMB* and *GZMH*. **e**, UMAP of liver MAIT cells coloured by the 12 identified clusters. **f**, Expression of MAIT1 and MAIT17 genes in liver MAIT cell clusters. Dot size indicates the percentage of cells expressing the gene and dot colour indicates the level of expression. **g**, UMAPs of liver MAIT cells coloured by the expression of *GNLY*, *HSPA1A*, *CCL4*, *CCL20*, *IFNG* and *XCL1*.

Liver MAIT cells comprised 12 clusters (Fig 4e, Extended Data Fig. 4d). Analogous to blood, these showed modest transcriptional differences (Extended Data Fig. 4e; Supplementary Table 3b). Seven out of 12 clusters had fewer than 10 markers with a FC > 1.5 and 25% of genes, including *CCL4* and *IFNG*, were markers for multiple clusters. Clusters did not correspond with known functionally discrete populations such as MAIT1 and MAIT17 (Fig. 4f). As in blood, there was a *GNLY*-expressing cluster (cluster 8) with low but statistically enriched *GZMB* expression (Fig. 4g; Extended Data Fig. 4e-g). Remaining clusters expressed different activation- or stress-induced molecules, including heat shock proteins, cytokines, and chemokines (Fig. 4g; Supplementary Table 3b). Although certain genes were predominantly expressed in a single cluster, most showed a gradient of expression across clusters.

Despite reported phenotypic, functional and/or transcriptional differences^6–8, 44^, CD8^+^, DN and CD4^+^ MAIT cells did not comprise separate clusters in blood or liver (Extended Data Fig. 4h-j). Moreover, the fractions of cells expressing different co-receptors were similar across clusters. CD8^+^ and DN MAIT cells differentially expressed two (*CD8A*/*B*) and eight genes in the blood and liver, respectively (Supplementary Table 4a-b). CD4^+^ and CD8^+^ MAIT cells differentially expressed 19 genes in blood and 43 in liver (Supplementary Table 4c-d), with 14 genes differentially expressed in both tissues. These included increased *CD4*, *CTSB* and *ITGB1* in CD4^+^ cells, and increased *CD8A*/*B*, *NKG7* and *PRF1* in CD8^+^ cells.

In summary, MAIT cells displayed tissue-specific gene expression and regulation, but functionally distinct subsets were not identified within tissues, with the potential exception of a *GNLY*-expressing cluster primed for cytotoxic activity.

### TCR clonotypes show variable bias in cluster localisation

As we had transcriptional and TCR data for each single cell, we examined whether limited transcriptional heterogeneity within tissues correlated with clonal origin. Given the donor-specific private TCRαβ repertoire we had identified, we performed clustering analysis separately for the seven samples. Multiple MAIT cell clusters were present in each sample, with clusters exhibiting limited transcriptional diversity (Extended Data Fig. 5; Supplementary Table 5). The median number of cluster marker per sample with FC > 1.5 was comparable to the multi-sample analyses (Extended Data Fig. 5h).

To statistically assess whether clonotypes were non-randomly distributed across clusters, we used the exact multinomial test. There was a range of associations between clonotype and cluster, both within and between different donors. Some TCR clonotypes were clearly skewed in distribution and predominantly localised in a single cluster (p < 0.01; Fig. 5a; Extended Data Fig. 5a). Other clonotypes showed a more subtle but still significant bias in cluster localisation (p < 0.01; Fig. 5b; Extended Data Fig. 5a). Some were randomly distributed (p = 1, Fig. 5c; Extended Data Fig. 5a). Bias in cluster distribution was more frequently significant for larger clonotypes (Fig. 5d), suggesting lack of significance for some smaller clonotypes may reflect lack of statistical power. For a given clonotype, the extent of bias in cluster localisation was not necessarily concordant in blood and liver (Fig. 5a,e). This could partly reflect the different size of the clonotypes in the two tissues. Nevertheless, some clonotypes displayed a stable transcriptional phenotype in blood and liver. For example, a *TRAV1-2*/*TRAJ12* clonotype (clonotype 3) from donor 2 preferentially localised to the *GNLY*-expressing cluster in blood (Fig. 5f,g) and liver (Fig. 5h,i). Therefore, the transcriptional profile of resting MAIT cells is influenced by their clonal origin.

**Fig. 5.**
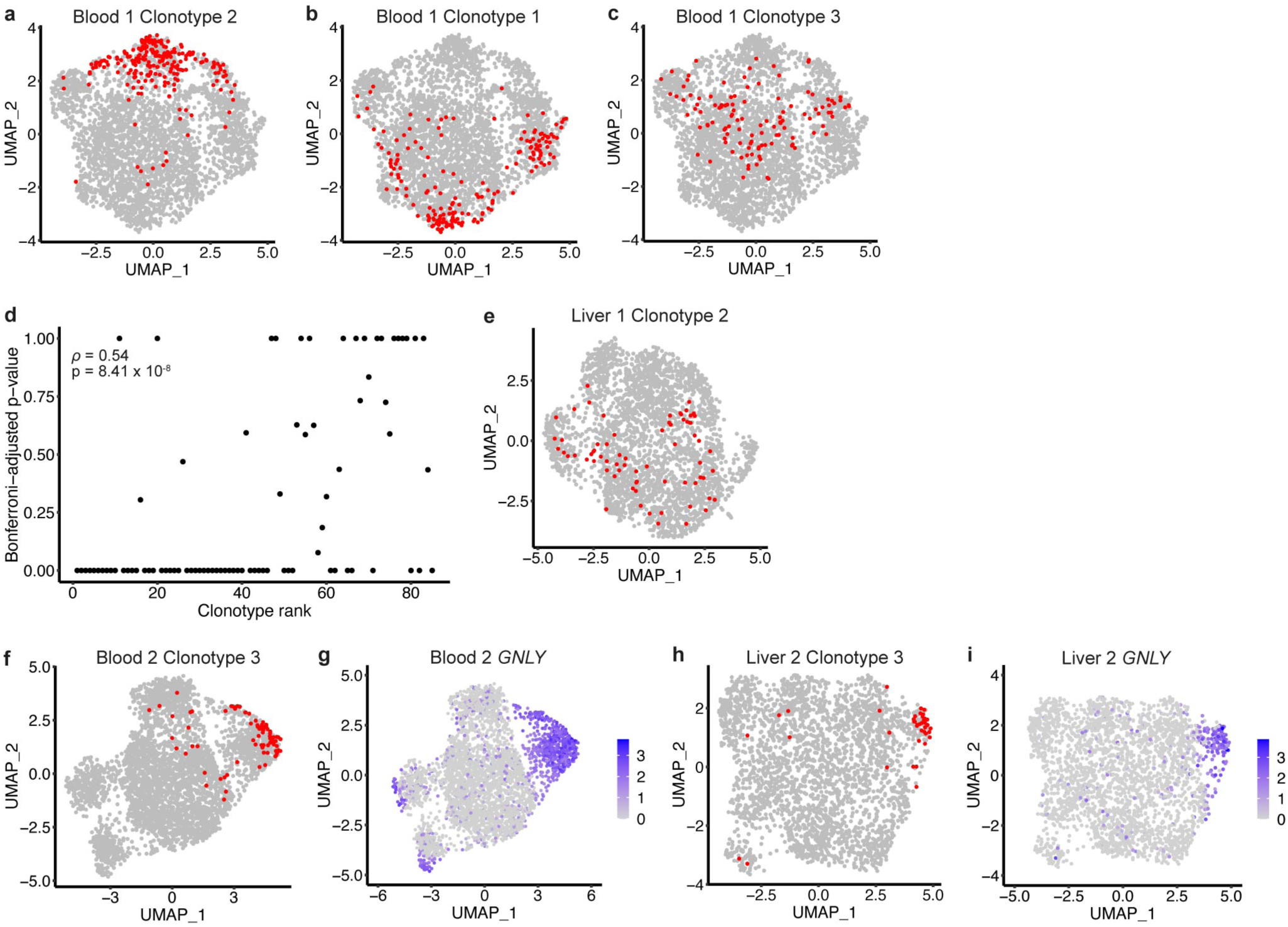
TCRαβ clonotypes are variably associated with transcriptional clusters. **a**-**c,** UMAPs of blood MAIT cells from donor 1 showing in red cells from a single TCRαβ clonotype. Plots show clonotype 2 (**a**), clonotype 1 (**b**) and clonotype 3 (**c**). Clonotype 1 is the largest clonotype from a donor, clonotype 2 is the second largest, and so on. **d**, Relationship between clonotype size (rank) and the Bonferroni-adjusted p-value for association between clonotype and cluster (exact multinomial test). Spearman’s rank correlation. **e**, UMAP of liver MAIT cells from donor 1 with cells from clonotype 2 indicated in red (same clonotype as in **a**). **f**,**g**, UMAP of blood MAIT cells from donor 2 with cells from clonotype 3 indicated in red (**f**) or expression of *GNLY* shown in purple (**g**). **h**,**i**, UMAP of liver MAIT cells from donor 2 with cells from clonotype 3 indicated in red (**h**) or expression of *GNLY* shown in purple (**i**).

### MAIT cell functional heterogeneity is governed by the nature of the activation stimulus

Given that we did not identify distinct subsets of human MAIT cells at baseline, we investigated whether functional subsets were present following activation. Isolated CD8^+^ T cells were left unstimulated or stimulated with plate-bound MR1/5-OP-RU or the cytokines IL-12 and IL-18. After 20 hours, MAIT cells were sorted (Extended Data Fig. 6) for scRNA-seq and scTCR-seq, an approach we termed functional RNA-seq (fRNA-seq). In total, 27,305 cells from the three donors and three experimental conditions passed quality control.

fRNA-seq revealed stimulus-specific transcriptional responses (Supplementary Table 6), with TCR- and cytokine-stimulated MAIT cells localising to different regions of the UMAP (Fig. 6a). We identified nine clusters of MAIT cells – these were present in all donors but were largely stimulus-specific (Fig 6b,c; Extended Data Fig. 7a). Consistent with a homogeneous resting transcriptional profile, unstimulated MAIT cells were predominantly (82%) found within a single cluster (cluster 0). TCR-stimulated cells were mostly in clusters 1 and 4. Cells in cluster 1 were more activated than those in cluster 4, displaying increased expression of chemokines and cytokines including *IFNG*, *TNF* and *CCL4* (Fig. 6d,e; Extended Data Fig. 7b; Supplementary Table 7). Clusters 2, 3 and 5 were largely comprised of cytokine-stimulated cells and appeared to reflect different levels of cell activation. Cells within cluster 3 expressed low levels of activation and effector molecules, such as *IFNG*, *IL2RA*, *GZMB* and *IL26*, while these genes were highly expressed in cluster 5 (Fig. 6d,e; Extended Data Fig. 7b; Supplementary Table 7). Cells in cluster 2 expressed high levels of *IFNG* but less *GZMB* than cells in cluster 5. Cells within cluster 7 (mostly cytokine-stimulated) uniquely expressed IFN-stimulated genes (Fig. 6d; Extended Data Fig. 7b).

**Fig. 6.**
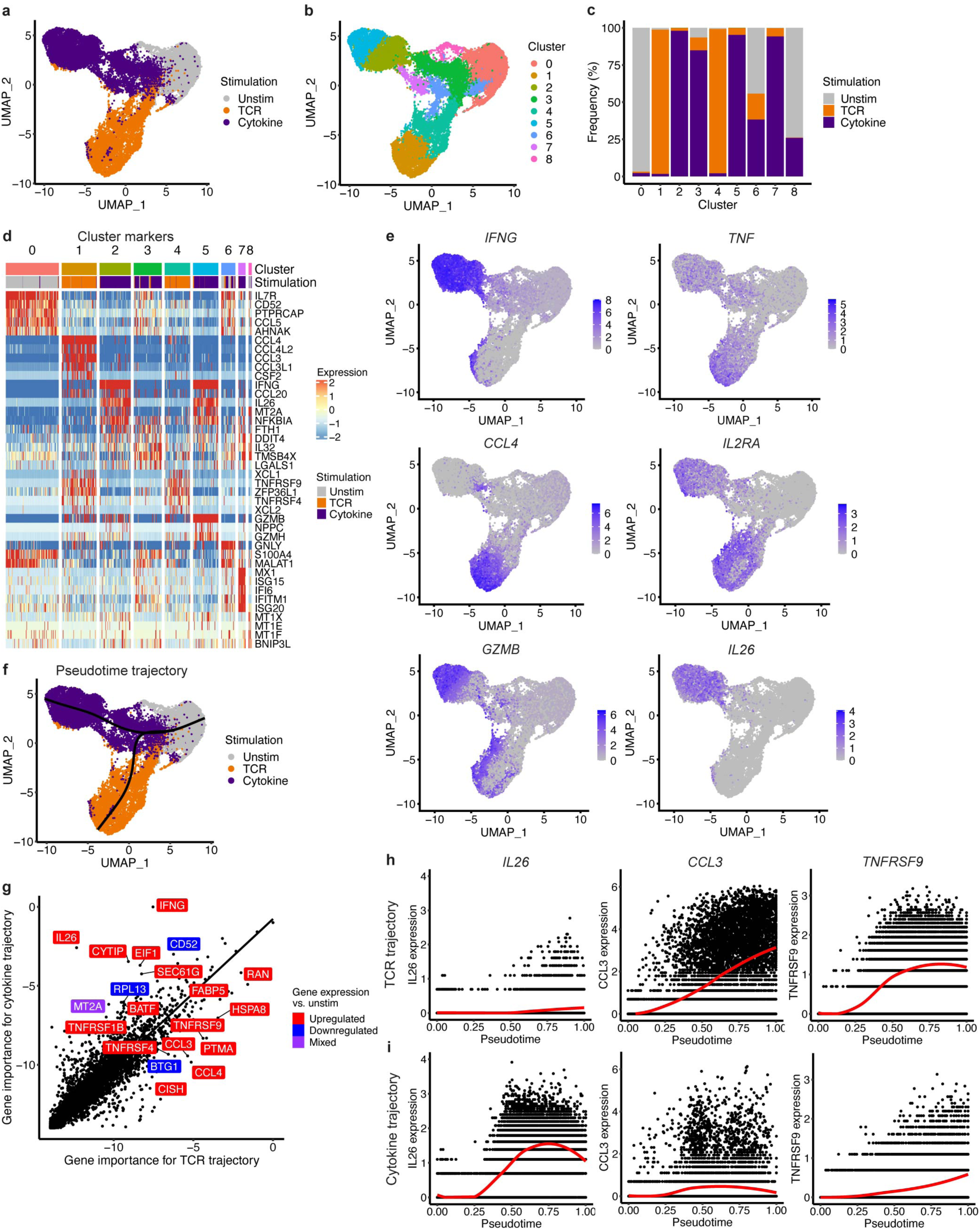
TCR- and cytokine-activated MAIT cells follow distinct linear trajectories. **a**,**b**, UMAP of MAIT cells from all donors coloured by stimulation condition (**a**) or the nine identified clusters (**b**). **c**, Proportion of cells in each cluster from the three stimulation conditions. **d**, Heatmap showing row-scaled log-transformed normalised expression of the top 5 marker genes for each cluster. **e**, UMAPs coloured by the expression of *IFNG*, *TNF*, *CCL4*, *IL2RA*, *GZMB* and *IL26*. **f**, UMAP showing in black the branching pseudotime trajectory identified using Slingshot. **g**, Spearman’s rank correlation of gene importance (log_2_ 1/gene importance rank) between the SCORPIUS TCR and cytokine trajectories. Labels show the most differentially important genes, 10 with higher importance on the TCR trajectory and 10 with higher importance on the cytokine trajectory. Colours indicate whether gene expression was upregulated (red), downregulated (blue) or mixed (purple; upregulated in TCR and downregulated in cytokine, or vice versa) relative to unstimulated cells. **h**,**i**, Expression of *IL26*, *CCL3* and *TNFRSF9* along the SCORPIUS TCR (**h**) and cytokine (**i**) trajectories.

As in the first experiment, we identified a distinct *GNLY*-expressing cluster (cluster 6; Fig 6d; Extended Data Fig. 7c). Strikingly, cluster 6 comprised cells from all three experimental conditions (Fig. 6c). These cells did not express other cytotoxic molecules such as *GZMB* or markers of activation, inconsistent with our initial hypothesis that *GNLY*-expressing cells at rest are primed for cytotoxic activity (Fig. 4b,d).

It has been suggested that expression of CD56 and/or CD94 identifies a subset of MAIT cells with enhanced cytokine responsiveness^9^. Following cytokine stimulation, CD56 (*KLRD1*)- and CD94 (*NCAM1*)-expressing MAIT cells showed increased IFNY production relative to their non-expressing counterparts (Extended Data Fig. 7d,e). However, *NCAM1* and *KLRD1* were expressed across multiple clusters, indicating that expression of these markers does not define a transcriptionally distinct cluster of MAIT cells (Extended Data Fig. 7f,g). Moreover, *KLRD1* was significantly upregulated following cytokine stimulation (Supplementary Table 6b; Extended Data Fig. 7g) and *NCAM1* appeared qualitatively increased (upregulation could not be formally tested due to a low gene detection rate; Extended Data Fig. 7f). Thus, it is difficult to assess the usefulness of these markers as baseline indicators of enhanced functional potential.

### Pseudotime analysis supports linear trajectories of MAIT cell activation

As MAIT cell clusters appeared to capture cells at different stages of activation, we further explored transcriptional responses to TCR and cytokine stimulation through pseudotime analysis. In accordance with the shape of the UMAP, the Slingshot^45^ pseudotime algorithm identified a branching trajectory with a single branch point close to unstimulated cells (Fig. 6f), suggesting MAIT cells become transcriptionally distinct early following TCR and cytokine stimulation. The presence of stimulus-specific trajectories starting from a homogeneous unstimulated population and transitioning to highly activated MAIT cells through multiple intermediate activation states was confirmed using SCORPIUS^46, 47^, an alternative pseudotime algorithm (Extended Data Fig. 7h,i).

Using random forest regression, we identified the genes most important for predicting cell pseudotime along the TCR and cytokine trajectories (Supplementary Table 8). Gene importance was highly correlated across the two trajectories (Fig. 6g), with 9 of the top 20 genes overlapping, including *IL2RA*, *TNFRSF18* (GITR) and *PKM* (all upregulated), and *IL7R* (downregulated). However, several genes of interest were important primarily for a single trajectory (Fig. 6g-i). Namely, *IFNG* and *IL26* were specific to the cytokine trajectory, while *CCL3*, *CCL4*, and *TNFRSF9* (4-1BB) showed greater importance for the TCR trajectory.

### Transcription factor activity differs in TCR- and cytokine-stimulated MAIT cells

To examine the regulation of shared and stimulus-specific gene expression, we used pySCENIC. We identified 159 high confidence regulons. Changes in transcription factor activity relative to unstimulated cells were significantly correlated for TCR- and cytokine-stimulated cells (Fig. 7a). Nevertheless, there were regulons with markedly different activity between the two conditions.

**Fig. 7.**
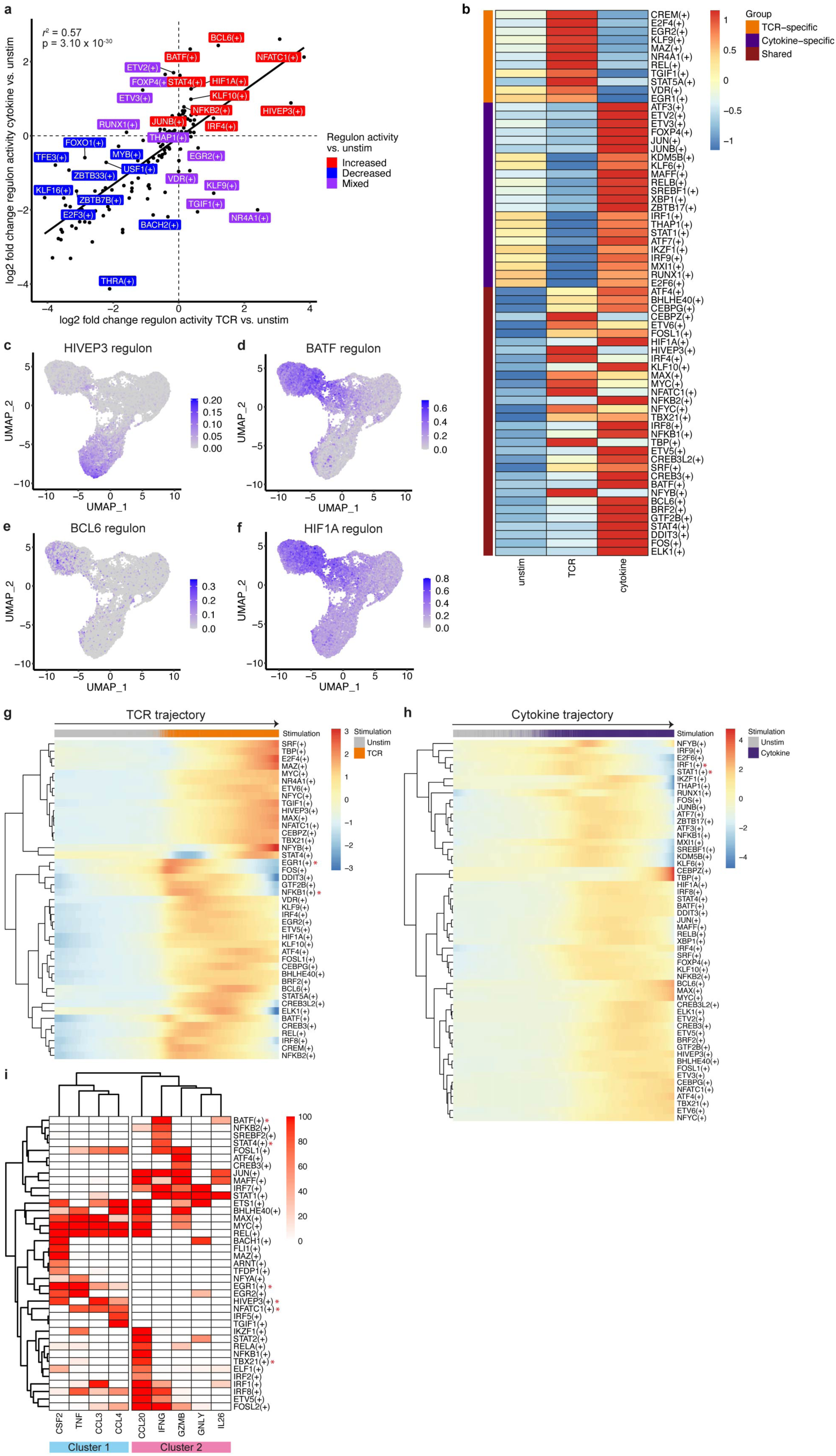
Transcriptional regulation of TCR- and cytokine-stimulated MAIT cells exhibits shared and distinct properties. **a**, Pearson correlation of the log_2_ fold change of regulon activity (AUCell scores) between TCR-stimulated and unstimulated MAIT cells, and cytokine-stimulated and unstimulated MAIT cells. Labels show the regulons with the largest difference in log_2_ fold change relative to unstimulated cells between the TCR and cytokine trajectory, 10 with increased (red), 10 with decreased (blue) and 10 with mixed (purple; increased in TCR and decreased in cytokine, or vice versa) activity following stimulation. **b**, Heatmap showing the activity (row-scaled average AUCell scores) of TCR-specific (orange), cytokine-specific (purple) and shared (maroon) upregulated regulons for each stimulation condition. **c**-**f**, UMAPs coloured by the activity of HIVEP3 (**c**), BATF (**d**), BCL6 (**e**) and HIF1A (**f**) regulons. **g**, Heatmap showing regulon activity (smoothed AUCell scores) over pseudotime on the SCORPIUS TCR trajectory for regulons upregulated upon TCR stimulation. **h**, Heatmap showing regulon activity (smoothed AUCell scores) over pseudotime on the SCORPIUS cytokine trajectory for regulons upregulated upon cytokine stimulation. **i**, Regulation of select MAIT cell effector genes. Heatmap is coloured by the percent occurrence of each gene within a given high confidence transcription factor regulon. Regulons included were predicted to regulate at least one of the genes in > 50% of pySCENIC runs. Red asterisks in **g**-**i** indicate regulons mentioned in the text. (+) in **a**, **b** and **g**-**i** indicates a regulon.

Focussing on the 65 regulons with significantly increased activity relative to unstimulated cells, 11 were TCR-specific, 22 were cytokine-specific and 32 were increased in both conditions (Fig. 7b). TCR-specific regulons included the TCR-induced transcription factors EGR1, EGR2 and NR4A1, as well as VDR, CREM and STAT5A, which have differing roles in the regulation of Th17 differentiation and IL-17 production^48–51^ (Fig. 7b; Extended Data Fig. 7j). Balance in their activity may regulate MAIT cell IL-17 production. Cytokine-specific regulons included STAT1, interferon regulatory factors (IRF1 and IRF9), XBP1 and IKZF1 (Ikaros) (Fig. 7b; Extended Data Fig. 7k). Ikaros regulates responsiveness to IL-12 signalling and IL-12Rα expression in conventional CD8^+^ T cells^52^ and may have a similar role in MAIT cells.

The 32 regulons with increased activity upon TCR and cytokine stimulation were also all significantly differentially active between the two conditions. TBX21 was most similar in activity between TCR- and cytokine-stimulated cells – its target genes were enriched for interleukin, Toll-like receptor and NF-kB signalling pathways (Extended Data Fig. 7l). As expected, NFATC1 and STAT4 showed enriched activity in TCR- and cytokine-stimulated cells, respectively. In addition, we identified novel candidate regulators of stimulus-specific MAIT cell functions. These included HIVEP3 for TCR-stimulated cells, and BATF, BCL6 and HIF1A for cytokine-stimulated cells (Fig. 7c-f). HIVEP3 is essential for the development of innate-like T cells including MAIT cells^53^. Amongst varied roles in T cell development and function, BATF promotes effector CD8^+^ T cell differentiation through upregulation of key transcription factors (e.g. T-bet), cytokine receptors (e.g. IL- 12 receptor) and signalling molecules^54^. Our data supports further work to investigate the role of these transcription factors in regulating MAIT cell effector functions.

As with upregulated genes, TCR- and cytokine-induced regulons showed a range of activity levels across stimulated cells. Most progressively increased in activity over pseudotime (Fig. 7g,h). However, activity of some regulons peaked early and subsequently declined, for example EGR1 and NFKB1 on the TCR trajectory, and STAT1 and IRF1 on the cytokine trajectory. Several early activated regulons regulated later activated transcription factors. As expected, STAT1 was a predicted regulator of TBX21^55^. HIVEP3 was a predicted target of EGR1, supporting its activation and function specifically in response to TCR signalling.

To identify candidate transcriptional regulators of MAIT cell effector genes, we examined their localisation within pySCENIC regulons. Unsupervised hierarchical clustering identified two clusters of effector genes (Fig. 7i). Cluster 1 comprised genes predominantly induced by TCR signalling, namely *CSF2*, *TNF*, *CCL3* and *CCL4* (Fig. 6g; Supplementary Table 6), suggesting similar regulation. Genes in cluster 1 were regulated by EGR1 and NFATC1. In addition, HIVEP3 was a predicted regulator of *CSF2*, *CCL3* and to a lesser extent *CCL4*. Cluster 2 comprised a mix of cytokine-specific genes, such as *IL26*, and genes induced by both stimuli, such as *CCL20* and *GZMB*. Surprisingly, IFNY was not present within the T-bet (TBX21) regulon but was regulated by STAT4 as expected. Additionally, IFNY was found within the BATF regulon, suggesting BATF could contribute to the enhanced IFNY production in cytokine- compared with TCR-stimulated MAIT cells.

Overall, TCR and cytokine stimulation induce shared and stimulus-specific changes in MAIT cell regulation. Our data suggest novel candidate regulators of TCR- and cytokine-specific responses and their predicted target genes, which can be tested experimentally.

### Clonotypic origin contributes to MAIT cell activation potential in response to both TCR and cytokine stimulation

General characteristics of the MAIT cell TCR repertoire in the fRNA-seq dataset were comparable to the blood-liver experiment (Extended Data Figure 8a-i). However, clonal diversity (reduced Gini coefficient, increased Shannon diversity index) was non-significantly increased, probably due to the younger age of the donors^37^ (blood-liver experiment: average 58, range 50-65 years; stimulation experiment: average 28, range 25-31 years) (Extended Data Fig. 8j,k). Similar to the blood-liver experiment, there was a trend towards increased use of *TRBV6-4* amongst *TRAJ12* and *TRAJ20* TCRs, however *TRAJ33* TCRs did not show increased use of *TRBV20-1* (Extended Data Fig. 8l). *TRAJ*- *TRBV* pairing differed considerably between donors, consistent with the blood-liver data (Extended Data Fig. 8m).

TCR clonotypes showed varied association with transcriptional clusters at rest (Fig. 5) and published data suggests functional differences related to TCRβ chain usage^9, 29, 30^. Using fRNA-seq, we investigated whether MAIT cell activation potential, as quantified by pseudotime position, was related to TCRβ chain usage or clonotypic origin. Within donors, activation capacity was significantly associated with *TRBV* usage, but there was clear variability amongst cells expressing the same *TRBV* gene segment (Fig. 8a,b). Results were consistent across two trajectory analysis methods (SCORPIUS and Slingshot) (Extended Data Fig. 9a,b). There was no correlation in *TRBV* activation potential between donors (Fig. 8a,b; Extended Data Fig. 9c,d) or between TCR and cytokine trajectories (Fig. 8c). Thus, there is no intrinsic difference in the activation potential of different *TRBV* gene segments.

**Fig. 8.**
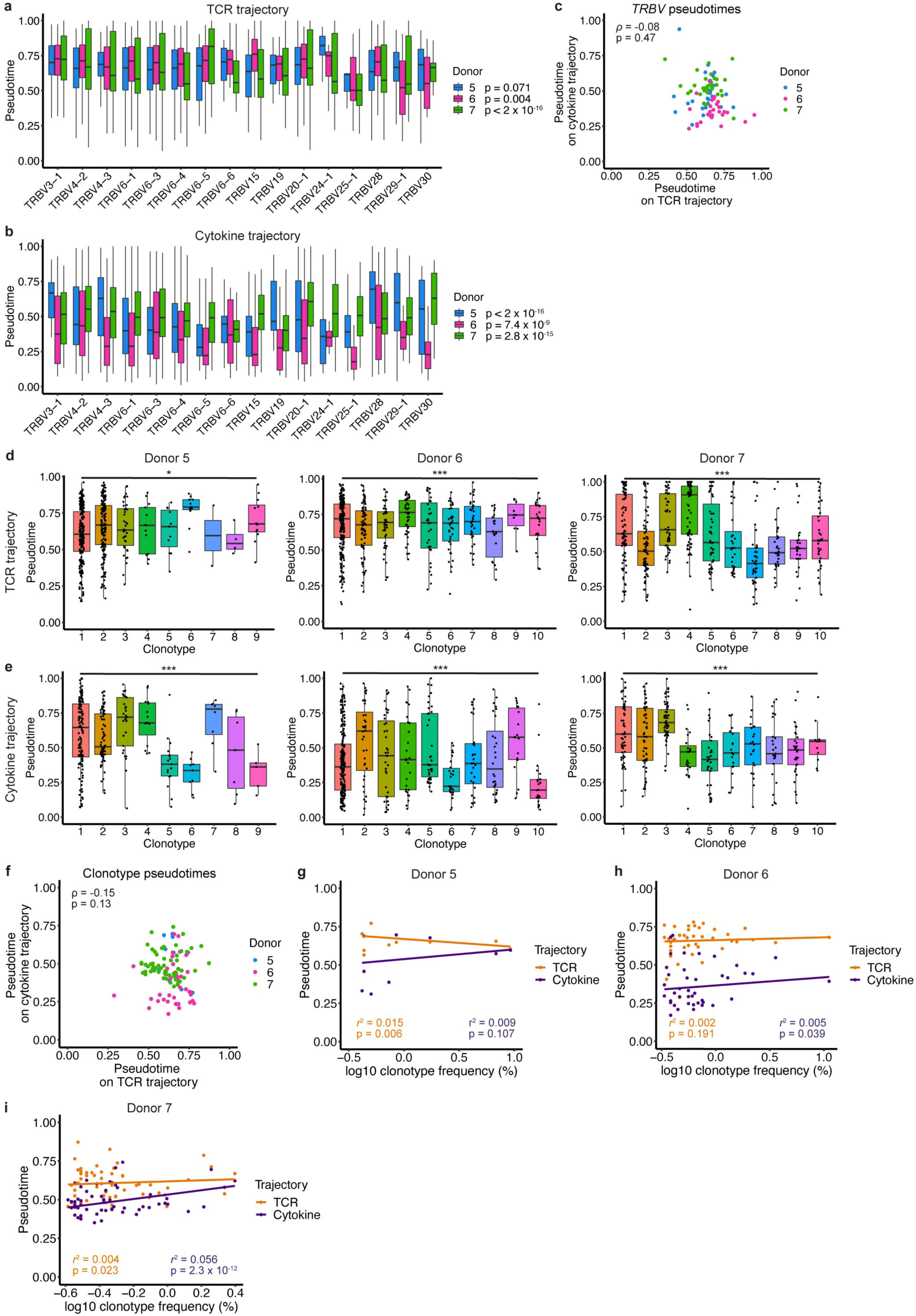
Clonotypic origin contributes to MAIT cell activation potential. **a**,**b**, Box plots split by donor showing pseudotime values on the SCORPIUS TCR (**a**) and cytokine (**b**) trajectories for MAIT cells expressing different *TRBV* gene segments. **c**, Correlation in average *TRBV* pseudotimes on the SCORPIUS TCR and cytokine trajectories. **d**,**e**, Pseudotime values for the largest 10 clonotypes per donor (or all clonotypes containing > 20 cells) on the SCORPIUS TCR (**d**) and cytokine (**e**) trajectories. **f**, Correlation in average clonotype pseudotimes on the SCORPIUS TCR and cytokine trajectories. **g**-**i**, Correlation between log_10_ clonotype frequency and pseudotime on the SCORPIUS TCR and cytokine trajectories for donor 5 (**g**), 6 (**h**) and 7 (**i**). Plots show stimulated cells only, *TRBV* gene segments with a frequency > 1% in any donor, and clonotypes containing > 20 cells. One-way ANOVA in **a**, **b**, **d**, **e**. One-way ANOVA in **a**, **b** performed for *TRBV* gene segments shown, and in **d**, **e** performed for all clonotypes containing > 20 cells. Spearman’s rank correlation in **c**, **f** and Pearson correlation in **g**-**i**. * p < 0.05, *** p < 0.001.

Based on these data, we hypothesised that observed variation between TCRs with different *TRBV* gene segments may reflect differences in the activation capacity of individual clonotypes. Consistent with this, activation capacity significantly differed between clonotypes on the TCR and cytokine trajectory (Fig. 8d,e), with high correlation between the two trajectory analysis methods (Extended Data Fig. 9e,f). High variability within clonotypes (Fig. 8d,e) suggested additional major influences on MAIT cell activation capacity. Activation capacity was not correlated in response to TCR and cytokine stimulation (Fig. 8f), and there was no consistent association between clonotype size and responsiveness to stimulation (Fig. 8g-i). Therefore, our data does not support the hypothesis that larger clones have expanded relative to other clones due to higher intrinsic functionality. Given that variation between clonotypes was observed on the cytokine as well as the TCR trajectory, differences in clonotype functionality appear to be cell-intrinsic rather than directly linked to the affinity of TCR-ligand binding. In summary, clonotypic origin contributes to the activation potential of an individual MAIT cell in response to both TCR and cytokine stimulation, although the factors that govern this relationship remain to be fully elucidated.

## Discussion

In summary, our single-cell data from human blood and liver, and from TCR- and cytokine-stimulated cells, suggests that human MAIT cells comprise a single functional population. Transcriptional plasticity is governed by tissue localisation, clonotypic origin and activation state. Despite their semi-invariant TCR, diverse TCRβ usage results in private MAIT cell TCR repertoires. The clonal identity of an individual cell influences both its resting and activated transcriptional profile.

Liver MAIT cells were transcriptionally distinct from blood, exhibiting an activated and tissue-resident gene expression profile. Most transcriptional changes were observed throughout the MAIT cell population, supporting a model whereby MAIT cells undergo environmentally driven adaptation following tissue entry. Basal activation of liver MAIT cells could reflect responses to microbial ligands transported from the gut to the liver via the hepatic portal vein^56^. Residency in the liver is consistent with bulk RNA-seq data from human liver MAIT cells and with mouse parabiosis experiments^12^. However, TCR repertoires were highly shared between matched blood and liver. MAIT cells are present in human thoracic duct lymph at a frequency comparable to matched blood and TCR usage is highly overlapping in the two sites^25^. Moreover, in human intestinal and uterine transplantation, small intestinal and endometrial MAIT cells are largely recipient-derived when measured at 12- and 23/34-months post-transplantation, respectively^57, 58^. Therefore, the extent of human MAIT cell tissue residency requires further examination. Nevertheless, our study adds to the growing body of literature demonstrating adaptation of human MAIT cell phenotype and function to tissue localisation^12, 18–20, 27, 58–60^.

Within tissues, transcriptional heterogeneity was low. Observed diversity indicated different activation states rather than distinct lineages. In contrast, mouse MAIT cells comprise developmentally, transcriptionally, and functionally distinct MAIT1 and MAIT17 subsets^10, 11, 41, 42^. The MAIT cell TCR and the antigen-presenting molecule MR1 are highly evolutionarily conserved^40, 61–63^. Thus, it is unclear why MAIT cells have evolved to include multiple specialised subsets in mice and one multifunctional population in humans, or the extent to which species disparities reflect different degrees of microbial exposure. Despite prior evidence of differences in phenotype and function dependent on co-receptor expression^6–8, 44^, human CD4^+^, CD8^+^ and DN MAIT cells did not form distinct clusters at a single-cell level (Extended Data Fig. 4h-j), as previously shown for mouse MAIT cells^64^. This indicates relatively minor transcriptional differences, but should be confirmed with larger numbers of CD4^+^ MAIT cells and in a setting where DN MAIT cells can be positively identified at the protein level, for example using hashing antibodies with scRNA-seq.

Despite modest diversity at rest, the MAIT cell transcriptome was highly plastic upon activation, with cells falling along stimulus-specific activation trajectories. Transcriptional plasticity is consistent with prior studies showing that MAIT cells produce high levels of type 17 cytokines^24^ and may even acquire type 2 functionality^23^ following prolonged stimulation. Transcriptional responses to stimulation were underpinned by alterations in regulatory networks. Although many genes and transcription factors showed similar expression and regulatory activity, respectively, along the TCR and cytokine trajectories, we identified stimulus-specific genes and regulators. For example, *CCL3* and *CCL4* were more strongly upregulated by TCR stimulation and were predicted to be regulated by HIVEP3, a TCR-specific transcriptional regulator (Fig. 6g; Fig. 7a,i). *IFNG* and *IL26* were more strongly upregulated by cytokine stimulation and were predicted to be regulated by BATF, a cytokine-specific transcriptional regulator (Fig. 6g; Fig. 7a,i). Roles for HIVEP3 and BATF in regulating MAIT cell functional responses to TCR and cytokine stimulation, respectively, have not previously been described. The cytokine-specific activity of BATF and its increased activity in liver suggests liver MAIT cells may be receiving tonic cytokine signals. Likewise, increased EGR1 activity in the liver suggests tonic TCR signalling.

It has been proposed that the MAIT cell response to cytokine stimulation is more restricted than the response to TCR stimulation^22^. Targeted studies, including standard flow cytometry approaches, are not well suited to accurately profile the MAIT cell functional response, due to the limited number of effector molecules examined, typically IFNY, TNF, IL-17 and granzyme B. Our genome-wide transcriptional data suggests equally broad but functionally distinct responses to TCR and cytokine stimulation. This finding should be validated at multiple timepoints, since the TCR response may peak earlier than the cytokine response^15, 22^. Nevertheless, our data should be considered in future analyses of cytokine-driven MAIT cell activation.

In resting blood and liver, as well as in activated MAIT cells, we identified a cluster of cells largely defined by their expression of *GNLY*. The proportion of *GNLY*-expressing cells in blood was comparable to previous measurements of *GNLY* protein^44^. Consistent with their unique cytotoxic profile^43, 44^, *GZMB* was lowly expressed in resting MAIT cells but was relatively enriched in the *GNLY*-expressing cluster in the blood and liver. This suggested that *GNLY*-expressing MAIT cells might be primed for cytotoxic activity. However, following TCR and cytokine stimulation, activated MAIT cells expressing high levels of *GZMB* did not co-express *GNLY*, and most *GNLY*-expressing cells from the three stimulation conditions were found within a single cluster close to unstimulated cells on the UMAP. Therefore, the function of this *GNLY*-expressing MAIT cell population requires further investigation.

Basic characteristics of the TCR repertoire were consistent with prior studies^8, 26, 35, 36, 40^. We identified a bias in TCRαβ pairing – *TRAJ12* and *TRAJ20* TCRα chains preferentially paired with *TRBV6-4* TCRβ chains, and *TRAJ33* TCRα chains with *TRBV20* TCRβ chains (Fig. 2j; Extended Data Fig. 3o,p). This could reflect an enhanced antigen affinity of these pairings resulting in increased microbe-mediated expansion in early life. However, our findings should be verified in a larger cohort given the modest effect size and wide variation between donors.

The extent of MAIT cell clonality was comparable to T_mem_ cells, with individuals displaying private TCRαβ repertoires. This challenges the paradigm of MAIT cells as a clonally restricted population with large numbers of public TCRs and corroborates previous studies that examined small cell numbers or bulk TCR repertoire data^37, 65^. However, the TCRα chain, which is key for ligand recognition^38, 39, 66^, was highly shared between individuals. Overlap of TCRαβ clonotypes in blood and liver was high. While consistent with shared TCRβ usage in matched blood and lymph^25^, this contrasts with other data indicating differential *TRAJ* or *TRBV* usage in tissues relative to blood, including breast^27^, kidney and intestine^26^. Although the degree of TCR sharing between blood and different tissues may vary, these studies lacked matched blood and tissue. Our study highlights the importance of matched samples when comparing TCR usage across tissues.

In resting blood and liver, we identified a novel association between the clonal origin and transcriptome of individual MAIT cells. This varied in strength between clonotypes and samples. Following stimulation, activation state significantly differed between clonotypes, despite wide variation within clonotypes. Clonal differences were observed following TCR and cytokine stimulation, but activation state did not correlate across the two trajectories. Our findings suggest intrinsic, stimulus-specific, and at least partly TCR-independent differences in the activation potential of MAIT cell clones. Based on our stimulation data, the cluster-clonotype association observed in resting blood and liver likely reflects differences in the activation state of clones/clusters. Altered activation capacity dependent on TCR clonotype is consistent with the altered clonal distribution following human *Salmonella* infection^29^ and increased MAIT cell clonality with age^37^. However, activation capacity was not correlated with clonotype size. This appears to contrast with a prior report demonstrating an enhanced proliferative capacity of MAIT cells expressing the most abundant Vβ segments following in vitro *Escherichia coli* stimulation^9^. Discordant results may be explained by considerable differences in the experimental approach. Overall, further work is necessary to understand the driving factors and functional consequences of clonal differences in activation capacity. While effect sizes were small, differences may be larger in the presence of suboptimal stimuli, such as weakly activating ligands or low concentrations of cytokines.

Our study provides improved understanding of human MAIT cell transcriptional, functional and TCR repertoire diversity, but we acknowledge several limitations. Due to cost restrictions, we were only able to study a small number of donors, two stimuli and a single time point following stimulation. However, large numbers of cells were included which was essential for establishing the extent of MAIT cell functional and TCR repertoire heterogeneity. Whilst our liver data is consistent with prior studies, it is unfeasible to obtain liver from healthy donors – uninvolved tissue from patients with liver tumours was used in the current study. Therefore, the described liver MAIT cell transcriptome may not be completely consistent with liver taken from a healthy donor. As previous studies showed altered kinetics of effector molecule production following TCR- and cytokine-dependent MAIT cell activation^15, 22^, some of our findings may be time-dependent. Analysis of transcription factor regulons gives insight into the regulation of gene expression, but high confidence regulons for some relevant transcription factors such as *PLZF* and *RORC* were not identified, perhaps due to poor gene detection.

In conclusion, we provide a genome-wide single-cell characterisation of the transcriptional profile and TCR repertoire of human blood and liver, and resting and activated, MAIT cells. Our data indicate private TCR repertoires with a high degree of sharing between blood and liver within an individual donor. MAIT cells showed stimulus-specific transcriptional responses, and we identified HIVEP3 and BATF as candidate regulators of the TCR- and cytokine-specific response, respectively. Although functionally distinct subsets were not identified at rest or following stimulation, clonality was variably linked with MAIT cell phenotype and function. Our data provide novel insights into human MAIT cell biology – relevant to other related innate-like subsets – and a comprehensive resource for further MAIT cell studies in health and disease.

## Acknowledgements

We thank Translational Gastroenterology Unit biobankers and Oxford University Hospitals NHS Foundation Trust surgeons for collection of patient blood and liver samples; S. Slevin for assistance with liver processing; K. Lynch for assistance with liver patient identification and sample processing; M. Salio for the plate-bound MR1 stimulation protocol; H. Ferry for sorting; D. Sims and D. Agarwal for discussions relating to computational analysis; and the NIH Tetramer Core Facility for MR1 monomers. Analysis was performed using computer systems at the MRC WIMM Centre for Computational Biology. L.C.G. is supported by a Wellcome PhD Studentship (109028/Z/15/Z). A.A. is supported by a Wellcome Clinical Training Fellowship (216417/Z/19/Z). M.E.B.F. is supported by Beyond Celiac and the Academy of Medical Sciences. N.M.P. is supported by an Oxford-UCB Postdoctoral Fellowship. P.K. is supported by the Wellcome Trust (WT109965MA), the NIH (U19 I082360), the NIHR Oxford Biomedical Research Centre and an NIHR Senior Fellowship.

## Author contributions

L.C.G., N.P. and P.K. designed the project and the experiments. L.C.G., A.A., M.E.B.F. and N.P. performed the experiments. L.C.G. performed the analysis. L.C.G. wrote the manuscript and all authors contributed to editing of the manuscript.

## Methods

### Data generation

#### Medium/buffers

R10: RPMI 1640 (Sigma-Aldrich), 10% FCS (Sigma-Aldrich), 1% Penicillin-Streptomycin (Thermo Fisher Scientific). Complete medium: R10, 1X non-essential amino acids (Thermo Fisher Scientific), 1 mM Na Pyruvate (Thermo Fisher Scientific), 10 mM HEPES (Thermo Fisher Scientific), 50 µM 2-mercaptoethanol (Thermo Fisher Scientific). FACS buffer: PBS, 0.5% BSA (Sigma-Aldrich), 1 mM EDTA (Sigma-Aldrich). Pre-sort buffer: PBS, 1% BSA, 10 mM HEPES.

#### Peripheral blood mononuclear cell (PBMC) isolation

PBMCs were isolated from fresh whole blood by density gradient centrifugation (Lymphoprep, Axis-Shield) at 2000 rpm (931 g) for 30 min with no brake. Cells were cryopreserved (90% FCS, 10% DMSO [Sigma-Aldrich]) in liquid nitrogen and thawed in complete medium as required on the day of use.

#### Liver tissue collection and processing

Liver tissue and matched blood for combined scRNA-seq and scTCR-seq were obtained from patients (50-65 years old) undergoing liver resection for metastatic colorectal cancer (n = 1; male), haemangioma (n = 1; female), or hepatocellular carcinoma (n = 2; one male and one female) at the Churchill Hospital, Oxford. Patients had no chronic liver disease, active excess alcohol consumption (> 14g/day), infection, immunosuppression, or family history of liver disease.

Disease-free liver tissue was collected from the resection margin and processed as previously described^67^. Briefly, liver tissue was cut into small pieces with a scalpel and ground through a 70 µm cell strainer. Cells were washed with R10 (2000 rpm [931 g], 10 min, 4°C) and mononuclear cells isolated by density gradient centrifugation on a discontinuous 35%/70% Percoll (GE Healthcare) gradient (2000 rpm [931 g], 20 min, room temperature, no brake). Mononuclear cells were collected from the interface and washed with R10 (1600 rpm [596 g], 10 min, 4°C). Residual red blood cells were lysed with ammonium-chloride-potassium (ACK) for 3-5 min. Cells were washed twice (1600 rpm [596 g], 10 min, 4°C) and cryopreserved (90% FCS, 10% DMSO) in liquid nitrogen.

#### Ethics

All samples were obtained with written informed consent through the Oxford Gastrointestinal Illnesses Biobank (REC Ref: 16/YH/0247).

#### In vitro stimulation of enriched CD8^+^ T cells

A Pierce streptavidin coated high capacity flat-bottom 96-well plate (Thermo Fisher Scientific) was coated with 50 µl biotinylated MR1/5-OP-RU monomer (NIH Tetramer Core Facility) at 10 µg/ml in PBS overnight at 4°C. Cryopreserved PBMCs from three healthy female donors were thawed in complete medium. CD8^+^ T cells were enriched using CD8 MicroBeads (Miltenyi Biotec) following manufacturer’s instructions. Enriched CD8^+^ T cells were washed in complete medium and resuspended at 1 x 10^7^ cells/ml.

One million cells per well were added to the appropriate 96-well plate (MR1/5-OP-RU-coated plate for the TCR stimulation condition, round-bottom plate for the unstimulated and cytokine stimulation conditions). IL-12 (50 ng/ml; R&D Systems) and IL-18 (50 ng/ml; R&D Systems) were added to the cytokine stimulation condition, αCD28 (1 µg/ml; clone: CD28.2; BioLegend) to the TCR stimulation condition, and complete medium to the unstimulated condition (final volume 200 µl in all wells). Cells were incubated for 20 hours at 37°C, 5% CO_2_.

#### Tetramer staining

Tetramer generation and staining was performed as previously described^68^ with minor modifications. Namely, biotinylated human MR1/5-OP-RU and MR1/6-FP monomers were provided by the NIH Tetramer Core Facility. Tetramers were generated using Streptavidin-Phycoerythrin or Streptavidin-Brilliant Violet 421 (BioLegend) following the NIH Tetramer Core Facility protocol. Tetramer staining was performed for 40 min at room temperature in FACS buffer.

#### Surface staining and fluorescence-activated cell sorting (FACS)

Surface staining was performed in Brilliant Stain Buffer (BD Biosciences) for 30 min at 4°C (reagents for tetramer and surface staining listed in Supplementary Table 9a,b). Cells were washed twice in PBS with 0.5% BSA, resuspended in pre-sort buffer containing 3-5 nM SYTOX Green Nucleic Acid Stain (Thermo Fisher Scientific), and incubated for 20 min at 4°C. Cells were sorted on a BD FACSAria III with a 70 µm nozzle. Sorted cells were collected in RPMI, 10% FBS, 25 mM HEPES. Sort purity was > 99%.

#### 10x Genomics library generation and sequencing

Sequencing libraries were generated using 10x Genomics Chromium Single Cell V(D)J Reagent Kits (v1.0 Chemistry) following manufacturer’s instructions. Cells were loaded onto the Chromium Controller (10x Genomics) at a concentration of ∼1 x 10^6^ cells/ml, with 6,000-8,000 cells loaded per channel. In most cases, one channel was loaded per donor. However, for donors 2 and 3 in the blood-liver experiment, sorted MAIT cells and T_mem_ cells were mixed at an equal concentration and loaded across two lanes of the Chromium Controller. Library quality and concentration was assessed using a TapeStation (Agilent) and Qubit 2.0 Fluorometer (Thermo Fisher Scientific), respectively. For the blood-liver experiment, libraries were sequenced on an Illumina HiSeq 4000 to a mean depth of 39,013-46,998 reads/cell for scRNA-seq (sequencing saturation 72.7-78.1%) and 10,182-30,495 reads/cell for scTCR-seq. For the stimulation experiment, libraries were sequenced on an Illumina NovaSeq 6000 to a mean depth of 76,000-89,000 reads/cell for scRNA-seq (sequencing saturation 70.0-90.1%) and 11,196-23,869 reads/cell for scTCR-seq. Library generation and sequencing were performed at the Oxford Genomics Centre (Wellcome Centre for Human Genetics, University of Oxford).

### Computational analysis

#### Pre-processing

10x Genomics Cell Ranger analysis pipelines (https://support.10xgenomics.com/single-cell-gene-expression/software) were used to generate single cell gene counts. For gene expression data, FASTQ files were generated from Illumina binary base call (BCL) files using cellranger mkfastq, then reads aligned to the 10x Genomics human GRCh38 reference genome (v3.0.0) and quantified using cellranger count (v3.0.1 for blood-liver experiment; v3.0.2 for stimulation experiment). For TCR sequencing data, BCL files were converted to FASTQ files using cellranger mkfastq (v2.1.1 for donors 1 and 4, v2.2.0 for donors 2 and 3 in blood-liver experiment; v3.0.2 for stimulation experiment) and single-cell V(D)J sequences generated using cellranger vdj (v3.0.1 for blood-liver experiment; v3.0.2 for stimulation experiment) with the 10x Genomics GRCh38 V(D)J reference (v2.0.0). The filtered contigs output file (filtered_contig_annotations.csv) was filtered to retain only high-confidence, full-length, productive contigs associated with TCRα or TCRβ chains. The data was then collapsed by cell barcode to generate a table (one per sample) with cells as rows and TCR regions as columns.

#### Quality control (individual samples)

Filtered feature-barcode matrices from cellranger count were imported into R and converted into a Seurat object (min.cells = 3, min.features = 200) using Seurat (v4.0.3)^69^. TCR and BCR genes were removed to ensure that downstream clustering analysis was not influenced by TCR or BCR chain usage.

For the blood-liver experiment, cells with very low or high unique molecular identifier (UMI) counts were removed. The lower UMI threshold, which aims to remove dead cells and cell-free RNA, was positioned at the local minimum of the UMI distribution leftwards of the mode UMI count. The upper UMI threshold was determined using a custom approach that makes use of matched scTCR-seq data to identify likely doublets (described previously^70^). Additional filtering was performed on gene counts and the percentage of mitochondrial reads – cells with ≤ 500 genes, ≥ 2,500 genes and/or ≥ 10% mitochondrial reads were removed. TCR chain information was added to the Seurat metadata slot. Cells with two TCRα and two TCRβ chains, or more than two TCRα and/or TCRβ chains, were assumed to be doublets and discarded. Cells with two TCRα and one TCRβ, or two TCRβ and one TCRα chain were retained, as a small fraction of T cells express two productive TCRα or TCRβ chains^71^.

For the stimulation experiment, cells with low UMI and gene counts (≤ 2 median absolute deviations below the median) and/or a high mitochondrial read fraction (≥ 2 median absolute deviations above the median) were removed. As UMI and gene counts were significantly increased upon TCR and cytokine stimulation, we chose not to apply an upper UMI threshold. Cells with two TCRα and two TCRβ chains, or more than two TCRα and/or TCRβ chains, were assumed to be doublets and discarded.

#### Normalisation, dimensionality reduction and clustering (individual samples)

Analysis was performed using Seurat^69^. Count data was normalised using sctransform^72^, with mitochondrial read fraction regressed out for samples in the blood-liver experiment. Highly variable genes were defined as the 3000 genes with the largest residual variance following variance stabilising transformation. Dimensionality reduction was performed using principal component analysis (PCA) with highly variable genes as input. Cell clusters were identified using Seurat’s graph-based clustering approach (0.5 resolution). Briefly, a shared nearest neighbour graph was built on dimensionally-reduced data (top 30 principal components [PCs]), then clusters determined by optimising the standard modularity function. Data was visualised by UMAP^73^ of the top 30 PCs. For blood-liver analyses, contaminating non-CD3^+^ T cell clusters (< 70% expressing *CD3E*) and non-MAIT cell clusters (< 70% expressing *TRAV1-2*), as well as clusters with low gene counts (significantly reduced [adjusted p < 0.01] compared with all other clusters as assessed by one-way ANOVA with Tukey’s honestly significant difference test), were removed before repeating the analysis steps from sctransform onwards. For samples containing MAIT and T_mem_ cells (donors 2 and 3), the two cell subsets were split following an initial clustering before analysis as described above.

#### Multi-sample analyses

For multi-sample analyses, a merged Seurat object (min.cells = 3, min.features = 200) was generated and filtered to retain only MAIT cells that passed quality control and filtering steps in individual sample analyses. TCR and BCR genes were removed. Normalisation and PCA were performed as in individual sample analyses with minor modifications. For the blood-liver experiment, highly variable genes were defined as the union of the top 1000 (all samples) or 1500 (blood, liver) highly variable genes from individual sample analyses. For the stimulation experiment, the top 3000 highly variable genes were directly defined for the merged sample. Samples were integrated using Harmony^74^ to remove batch effects between donors. Cell clusters were identified using the top 30 Harmony components with Seurat’s graph-based clustering approach (0.5 and 0.3 resolution for the blood-liver and stimulation experiment, respectively). Data was visualised by UMAP of the top 30 Harmony components.

#### Differential gene expression analysis

Differential gene expression analysis between clusters was performed using the Wilcoxon rank sum test for individual sample analyses and MAST^75^ (v1.18.0) for multi-sample analyses. Cluster markers were defined as genes with significantly increased expression in one cluster relative to the average of all other clusters (adjusted p < 0.05 based on Bonferroni correction using all genes in the dataset). Differential gene expression analysis between tissues, co-receptors and stimulation conditions was performed using MAST. Genes with a FC > 1.5 and a Bonferroni-adjusted p < 0.05 were defined as significantly differentially expressed. Analysis was performed using the FindMarkers function from Seurat. Input data was log-transformed normalised counts generated by global-scaling normalisation (NormalizeData function from Seurat). MAST analysis included UMI counts and donor as latent variables, or UMI counts only for the blood-liver comparison. For co-receptor analyses, cells with normalised expression values > 0 for *CD8A* and/or *CD8B* were defined as CD8^+^, cells with normalised expression values > 0 for *CD4* were defined as CD4^+^, and cells expressing neither *CD4*, *CD8A* or *CD8B* were defined as DN.

#### Pseudotime analysis

Pseudotime analysis was performed using Slingshot^45^ (v2.0.0) and SCORPIUS^46, 47^ (v1.0.8). UMAP coordinates and Seurat cluster labels (0.1 resolution) were provided as input to Slingshot, with the major unstimulated cluster specified as the start of the trajectory. Normalised expression values (sctransform) were provided as input to SCORPIUS. Two separate SCORPIUS trajectories were generated from unstimulated and TCR-stimulated cells, and unstimulated and cytokine-stimulated cells. Gene importance along SCORPIUS trajectories was determined using random forest regression (gene_importances function, num_permutations = 10). Differential gene importance was calculated by taking the ratio of gene importance ranks on the TCR and cytokine trajectories (higher rank as numerator). Genes defined as differentially important were required to have an importance false discovery rate (FDR) < 0.05 and to be within the top 150 most highly ranked genes for either the TCR or cytokine trajectory.

#### Transcription factor regulon analysis

Transcription factor regulons were identified using SCENIC^33, 34^ (pySCENIC v0.11.2). Briefly, the raw expression matrix was filtered to retain genes expressed in > 1% of cells and with a count > 3 x 0.01 x number of cells (612 for blood-liver, 819 for stimulation data). Modules comprising transcription factors and co-expressed genes were generated using GRNBoost2, then pruned to remove indirect targets lacking enrichment for the corresponding transcription factor motif (cisTarget). This resulted in a set of transcription factor regulons. Due to stochasticity in gene regulatory network inference using GRNBoost2, each pySCENIC run can identify a different number of regulons, as well as different target genes for each transcription factor. Thus, pySCENIC was run 100 times. High confidence regulons were defined as regulons that occurred in > 80% of runs and that contained at least five high confidence target genes. High confidence target genes were those found within a regulon in > 80% of runs. Cells were scored for the activity of each high confidence regulon (including only high confidence target genes) using AUCell (v1.14.0). Regulons differentially active between tissues or stimulation conditions were determined using MAST^75^ (FindMarkers function from Seurat) with donor as a latent variable. Regulons with a Bonferroni-adjusted p < 0.01 were defined as differentially active. Smoothed regulon activity scores (AUCell scores) over the SCORPIUS trajectories were generated by loess regression (loess function from the stats R package).

#### GSEA and pathway enrichment analysis

Pseudobulk gene counts were generated by summing raw gene counts for all cells within a sample. GSEA^76^ for published CD8^+^ T_RM_^31^ and CD4^+^ T_RM_ ^32^ cell gene signatures was performed using pseudobulk gene counts. Over-representation analysis for Gene Ontology (GO) terms or Reactome pathways was performed using the clusterProfiler R package (v4.0.0)^77^. Background genes were defined as all genes with a count ≥ 5 in the merged analysis of MAIT cells from all samples (seven and nine samples in the blood-liver and stimulation experiment, respectively). Redundant enriched GO terms were removed using the simplify function (measure = “Wang”).

#### Gene lists

MAIT1 and MAIT17 gene signatures were defined as the overlapping genes from two published scRNA-seq datasets^41, 42^. Mouse gene symbols were converted to human gene symbols using the getLDS function from the biomaRt R package (v2.48.1) with the Ensembl BioMart database and the “hsapiens_gene_ensembl” and “mmusculus_gene_ensembl” datasets. Interferon-stimulated genes were obtained from Schoggins and colleagues^78^.

#### Nucleotide and functional TCR clonotypes

Tables of TCRα and TCRβ usage for each cell in a sample (produced as described in pre-processing) were combined to generate one table per donor for nucleotide clonotype calling. For functional clonotype calling, a combined table was generated from all samples. TCR clonotypes were defined for cells that passed quality control and filtering steps in individual sample analyses. MAIT cells were additionally required to have a TRAV1-2 TCRα chain and at least one TCRβ chain. T_mem_ cells were required to have at least one TCRα and one TCRβ chain. Cells expressing TRAV1-2 paired with TRAJ33, TRAJ12 or TRAJ20 and with a 12 amino acid CDR3α region were assumed to be contaminating MAIT cells and removed before T_mem_ cell clonotype calling.

Nucleotide clonotypes were defined as cells with identical TCR gene segment usage, and CDR3α and CDR3β nucleotide sequences. TCRα only and TCRβ only clonotypes were defined as cells with identical TCRα segment usage and CDR3α sequences, or identical TCRβ segment usage and CDR3β sequences, respectively. TCRαβ clonotypes were numbered according to size, with clonotype 1 being the largest, clonotype 2 being the second largest, and so on. Clonotypes of identical size were randomly ordered for numbering. TCRαβ clonotypes were assigned ranks in a similar manner, but clonotypes of identical size were given the same rank.

Functional clonotypes were defined as cells with at least one identical TCRα and TCRβ chain amino acid sequence (TCR variable region gene segment usage and CDR3 amino acid sequences). Functional TCRα and functional TCRβ clonotypes were defined as cells with at least one matching TCRα or TCRβ chain amino acid sequence, respectively. Given the presence of TCR dropout, functional clonotypes were permitted to contain a mixture of cells with one or two TCRα or TCRβ chains, providing all detected chains matched those within the clonotype.

#### TCR analyses

TCR analyses were performed only for cells with a defined TCR clonotype. For the stimulation experiment, unstimulated cells were used for TCR repertoire characterisation. MAIT cell TCRα chain analysis was limited to the *TRAV1-2* TCRα chain. The Gini coefficient was calculated using the Gini function from the DescTools R package (v0.99.42). The Shannon diversity index was calculated using the diversity function from the vegan R package (v2.5.7). To test for an association between clonotype and cluster, the exact multinomial test was performed using the multinomial.test function from the EMT R package (v1.1; MonteCarlo = TRUE, ntrial = 1000000). *TRAJ*-*TRBV* pairings present in > 10 cells were included in Circos plots generated using the circlize R package (0.4.13). Sequence logos were generated using the ggseqlogo R package (v0.1). The overall height of the stacked letters at each position indicates the sequence conservation, while the relative abundance of each amino acid is indicated by the height of individual letters within the stack. Acidic bases are shown in red, basic residues in blue, hydrophobic in black and polar in green. The number of N and P mutations in CDR3α sequences of length 36 nucleotides were determined using IMGT/JunctionAnalysis^79^.

#### Plots

Most plots were generated using ggplot2 (v3.3.4). FACS plots were generated in FlowJo (v10.8.1). Heatmaps were generated using pheatmap (v1.0.12) or ComplexHeatmap (v2.8.0). GSEA plots were generated in Prism (v9.2.0).

## Tables#

• Supplementary Table 1 - differentially expressed genes between blood and liver MAIT cells

• Supplementary Table 2 – transcription factor regulons differentially active between blood and liver MAIT cells

• Supplementary Table 3a,b – blood cluster markers (a) and liver cluster markers (b)

• Supplementary Table 4a-d – differentially expressed genes between CD8+ and DN MAIT cells in the blood (a) and liver (b); differentially expressed genes between CD4+ and CD8+ MAIT cells in the blood (c) and liver (d)

• Supplementary Table 5a-g – cluster markers for individual sample analyses

• Supplementary Table 6a,b – differentially expressed genes between TCR-stimulated and unstimulated (a), and cytokine-stimulated and unstimulated (b), MAIT cells

• Supplementary Table 7 – cluster markers for the stimulation experiment

• Supplementary Table 8 – gene importance scores for SCORPIUS TCR and cytokine trajectories

• Supplementary Table 9a,b – reagents used for FACS in the blood-liver (a) and stimulation (b) experiments

---

**Extended Data Fig. 1.**
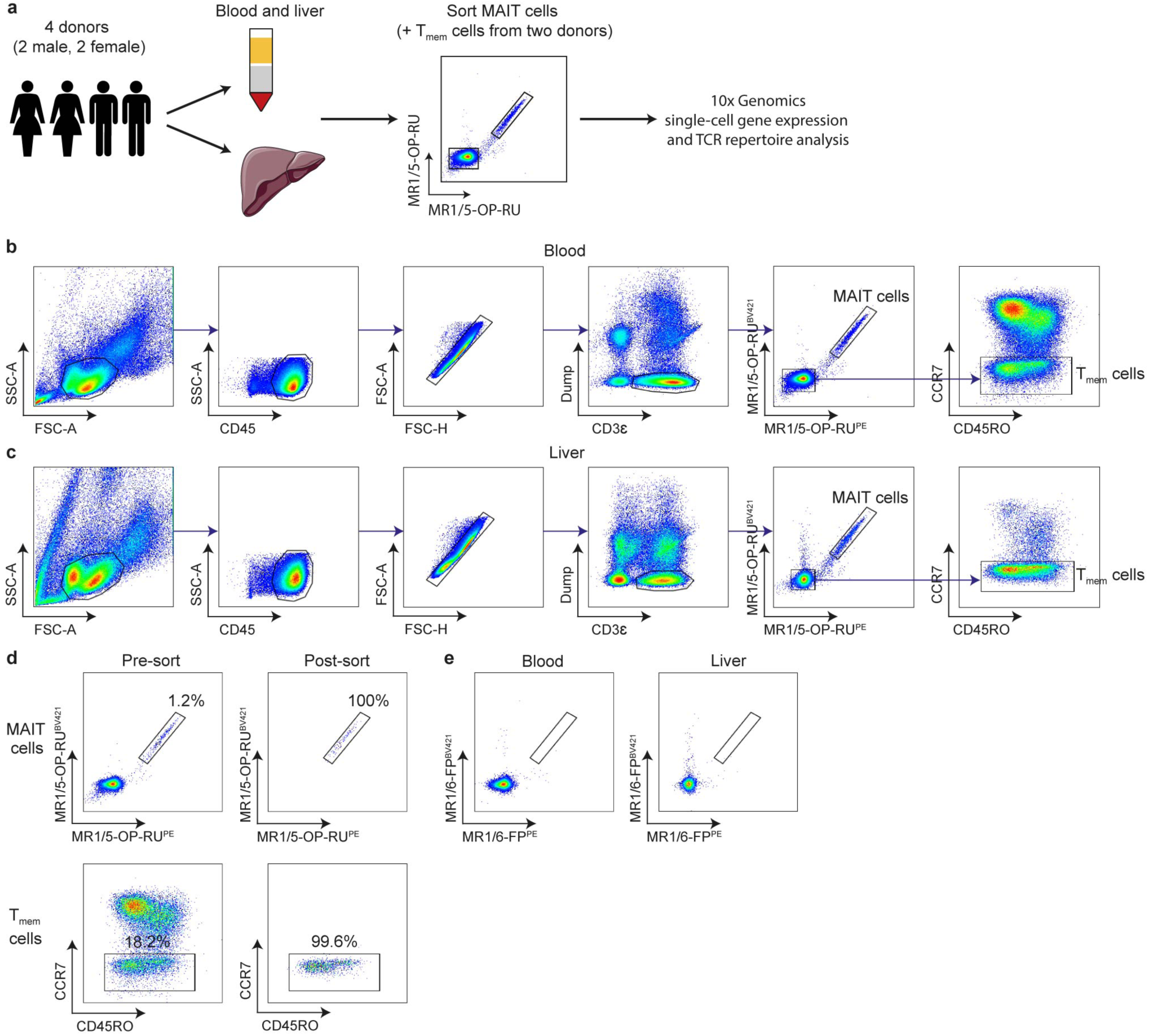
Blood-liver experiment schematic and gating strategy. **a**, Schematic illustrating the experimental protocol. **b**,**c**, Gating strategy for sorting MAIT cells (CD3^+^MR1/5-OP-RU^+^) and conventional memory T (T_mem_) cells (CD3^+^MR1/5-OP-RU^-^CCR7^-^) from matched human blood (**b**) and liver (**c**) for single-cell RNA-sequencing and single-cell TCR-sequencing. Dump channel contained antibodies to CD14, CD19, TCR γδ, TCR Vα24-Jα18 and TCR Vδ2, plus SYTOX Green Nucleic Acid Stain. **d**, FACS plots for a representative sample showing MAIT and T_mem_ cells pre- and post-sorting. Sort purity was > 99%. **e**, FACS plots for representative blood and liver samples showing staining with the negative control MR1/6-FP tetramer.

**Extended Data Fig. 2.**
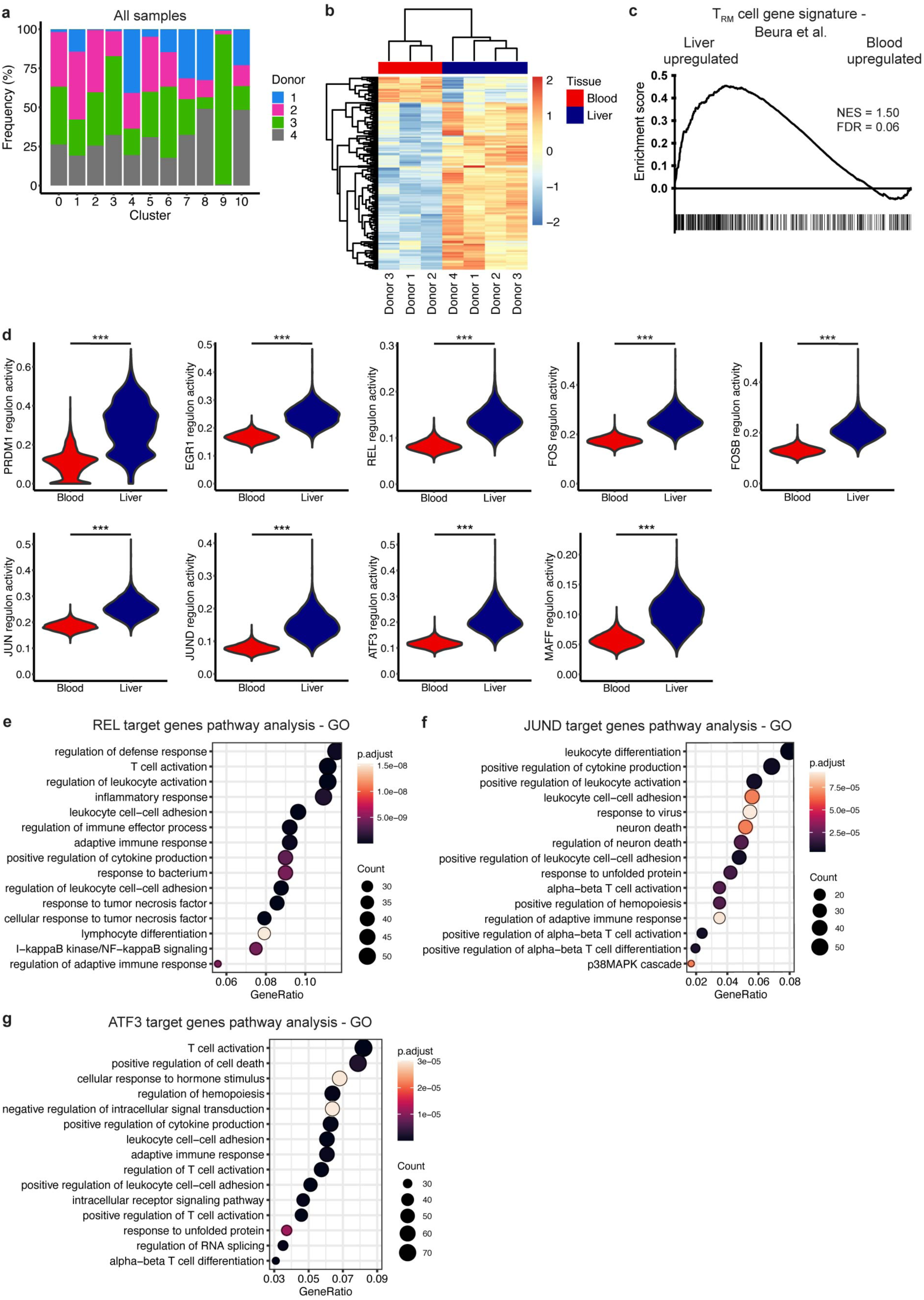
Blood and liver MAIT cells show distinct transcription and regulation. **a**, Proportion of MAIT cells from each donor in each cluster. **b**, Heatmap showing row-scaled log-transformed normalised (trimmed mean of M-values) pseudobulk expression of genes differentially expressed between blood and liver MAIT cells. **c**, Gene set enrichment analysis (GSEA) of liver compared with blood MAIT cells using a tissue-resident memory T (T_RM_) cell gene signature from Beura and colleagues^32^. NES = normalised enrichment score, FDR = false discovery rate. **d**, Violin plots showing the activity (AUCell scores) of selected liver-upregulated transcription factor regulons in blood and liver. Differential activity analysis was performed using MAST, *** p < 0.001. **e**-**g**, Pathway analysis on predicted REL (**e**), JUND (**f**) and ATF3 (**g**) target genes. Top 15 non-redundant gene ontology (GO) terms are shown.

**Extended Data Fig. 3.**
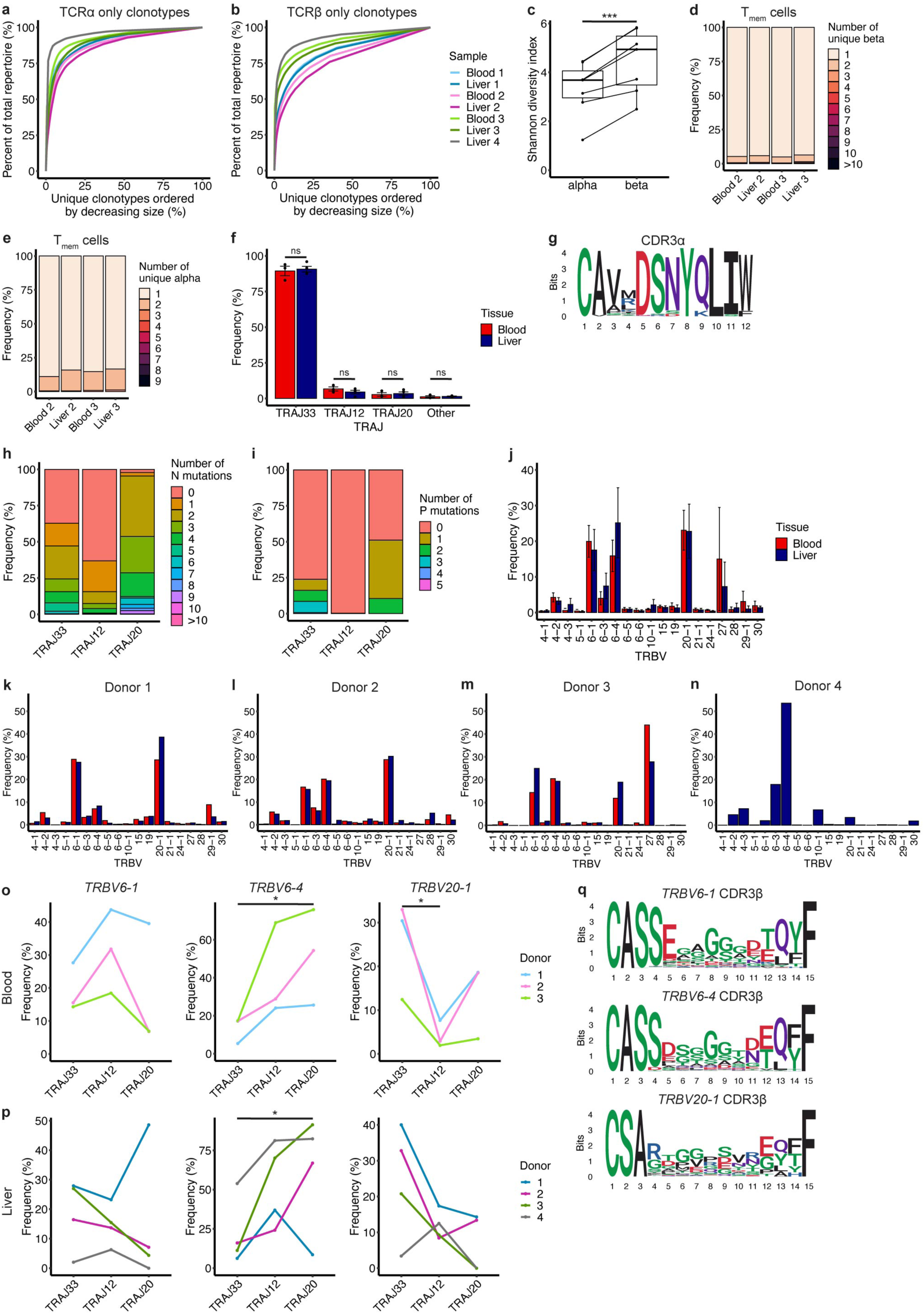
MAIT cells have a restricted TCRα but diverse TCRβ repertoire. **a**,**b**, Line plots demonstrating the clonality of the MAIT cell TCRα (**a**) and TCRβ (**b**) repertoire in each sample. **c**, Shannon diversity index for MAIT cell TCRα only and TCRβ only clonotypes. Paired t-test, *** p < 0.001. **d**,**e**, T_mem_ cell TCR chain pairing at the population level. Number of unique TCRβ chains paired with a single TCRα chain (**d**), or TCRα chains paired with a single TCRβ chain (**e**), in T_mem_ cells from each sample. **f**, Proportion of blood and liver MAIT cells expressing *TRAJ33*, *TRAJ12*, *TRAJ20* and other *TRAJ* gene segments. Mean ± SEM is shown. Unpaired t-test, ns = non-significant. **g**, Sequence logo generated from all MAIT cell CDR3α amino acid sequences of length 12. **h**,**i**, Number of N (**h**) and P (**i**) mutations in *TRAJ33*, *TRAJ12* and *TRAJ20* TCRα chains. Number of N and P mutations were significantly associated with *TRAJ* usage as measured by Fisher’s exact test (p < 0.0001 for both). **j**, Proportion of blood and liver MAIT cells expressing different *TRBV* gene segments. Plot includes *TRBV* gene segments present at a frequency > 1% in at least one donor. Mean ± SEM is shown. Unpaired t-test, blood-liver comparison non-significant for all *TRBV* gene segments. **k**-**n**, Proportion of blood and liver MAIT cells expressing different *TRBV* gene segments in donor 1 (**k**), 2 (**l**), 3 (**m**) and 4 (**n**). **o**,**p**, Percentage use of *TRBV6-1* (left), *TRBV6-4* (middle) and *TRBV20-1* (right) *TRBV* gene segments amongst *TRAJ33*, *TRAJ12* and *TRAJ20* MAIT cell TCRs in the blood (**o**) and liver (**p**). Friedman test with Dunn’s post-hoc test. Significant comparisons are indicated, * p < 0.05. **q**, Sequence logos generated from all MAIT cell *TRBV6-1*, *TRBV6-4* or *TRBV20-1* CDR3β amino acid sequences of length 15.

**Extended Data Fig. 4.**
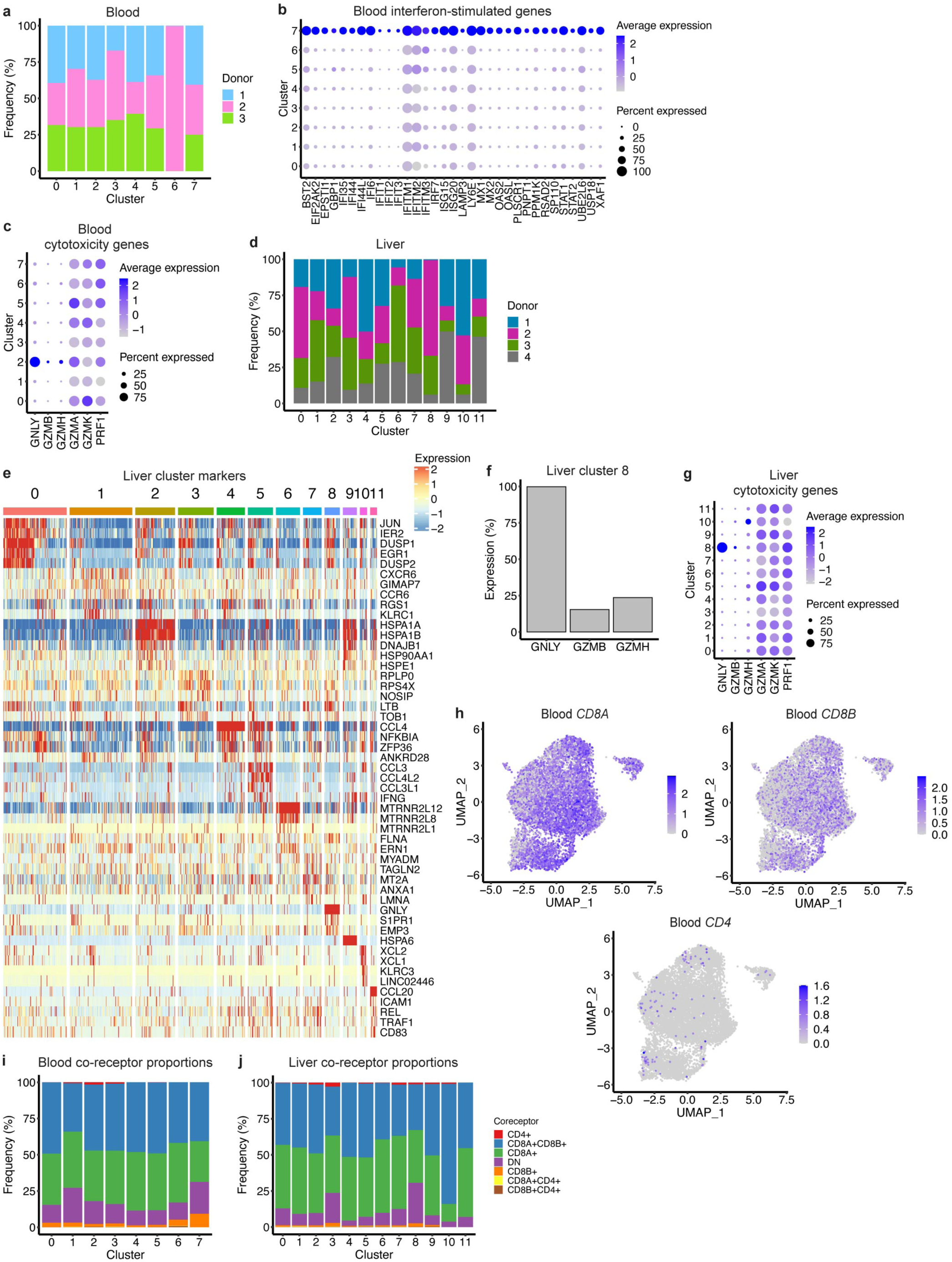
MAIT cells within the blood and liver show limited transcriptional heterogeneity. **a**, Proportion of MAIT cells from each donor in each blood cluster. **b**,**c**, Expression of select interferon-stimulated (**b**) and cytotoxicity (**c**) genes in blood MAIT cell clusters. Interferon-stimulated genes were markers for cluster 7. Dot size indicates the percentage of cells expressing the gene and dot colour indicates the level of expression. **d**, Proportion of MAIT cells from each donor in each liver cluster. **e**, Heatmap showing row-scaled log-transformed normalised expression of the top five marker genes for each liver MAIT cell cluster. **f**, Percentage of cells expressing (log-transformed normalised expression > 0) *GNLY*, *GZMB* and *GZMH* in liver cluster 8. **g**, Expression of select cytotoxicity genes in liver MAIT cell clusters. Dot size indicates the percentage of cells expressing the gene and dot colour indicates the level of expression. **h**, UMAPs of blood MAIT cells coloured by the expression of *CD8A*, *CD8B* and *CD4*. **i**,**j**, Proportion of MAIT cells in each blood (**i**) and liver (**j**) cluster expressing (log-transformed normalised expression > 0) the indicated co-receptor or combination of co-receptors.

**Extended Data Fig. 5.**
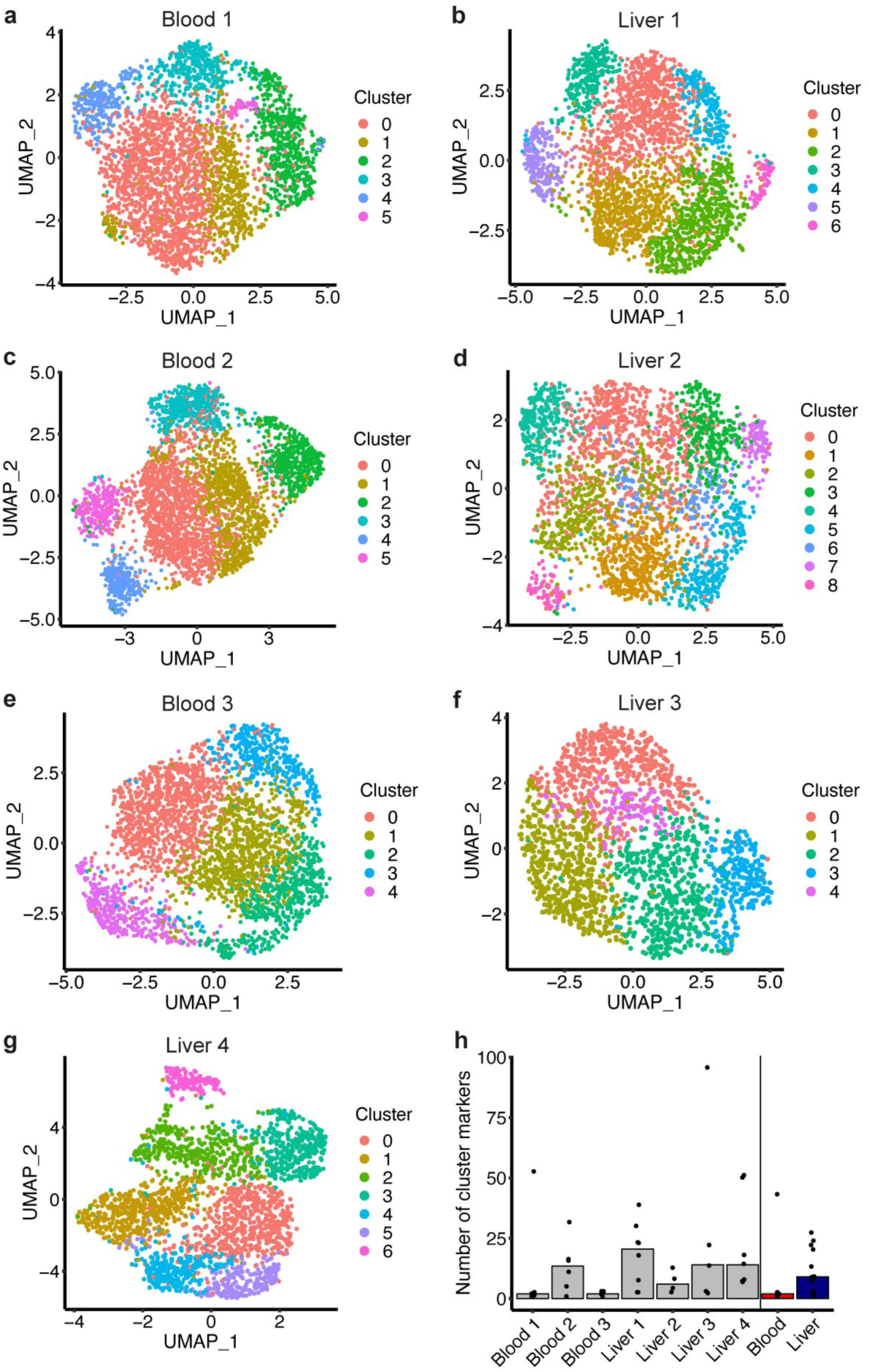
Clusters identified in individual samples. **a**-**g**, Individual sample UMAPs coloured by the identified clusters – blood 1 (**a**), liver 1 (**b**), blood 2 (**c**), liver 2 (**d**), blood 3 (**e**), liver 3 (**f**) and liver 4 (**g**). **h**, Median number of markers per cluster with fold change > 1.5 for individual sample (grey) and for merged blood (red) and liver (blue) analyses. Dots indicate the number of markers for individual clusters.

**Extended Data Fig. 6.**
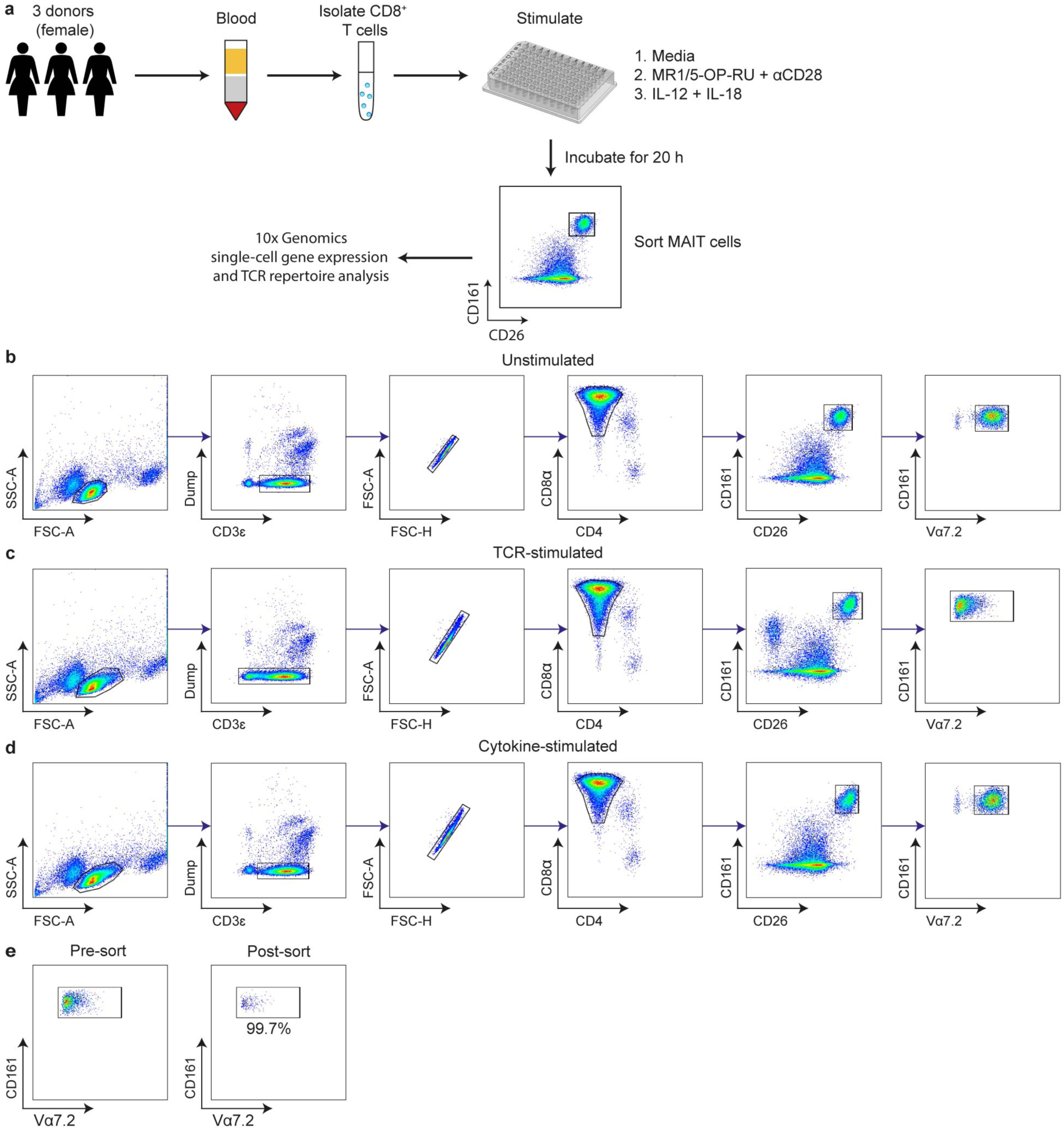
Stimulation experiment schematic and gating strategy. **a**, Schematic illustrating the experimental protocol. **b**-**d**, Gating strategy for sorting MAIT cells (CD3^+^CD8^+^CD26^+^CD161^hi^Vα7.2^+^) from unstimulated (**b**), TCR-stimulated (**c**) and cytokine-stimulated (**d**) enriched CD8^+^ cells for single-cell RNA-sequencing and single-cell TCR-sequencing. Due to TCR downregulation within the TCR stimulation condition (**c**), MAIT cells were sorted as CD8^+^CD26^+^CD161^hi^ lymphocytes. Dump channel contained antibodies to CD14, CD19, TCR γδ, TCR Vα24-Jα18 and TCR Vδ2, plus SYTOX Green Nucleic Acid Stain. **e**, FACS plots for a representative sample (TCR stimulation condition) showing MAIT cells pre- and post-sorting. Sort purity was > 99%.

**Extended Data Fig. 7.**
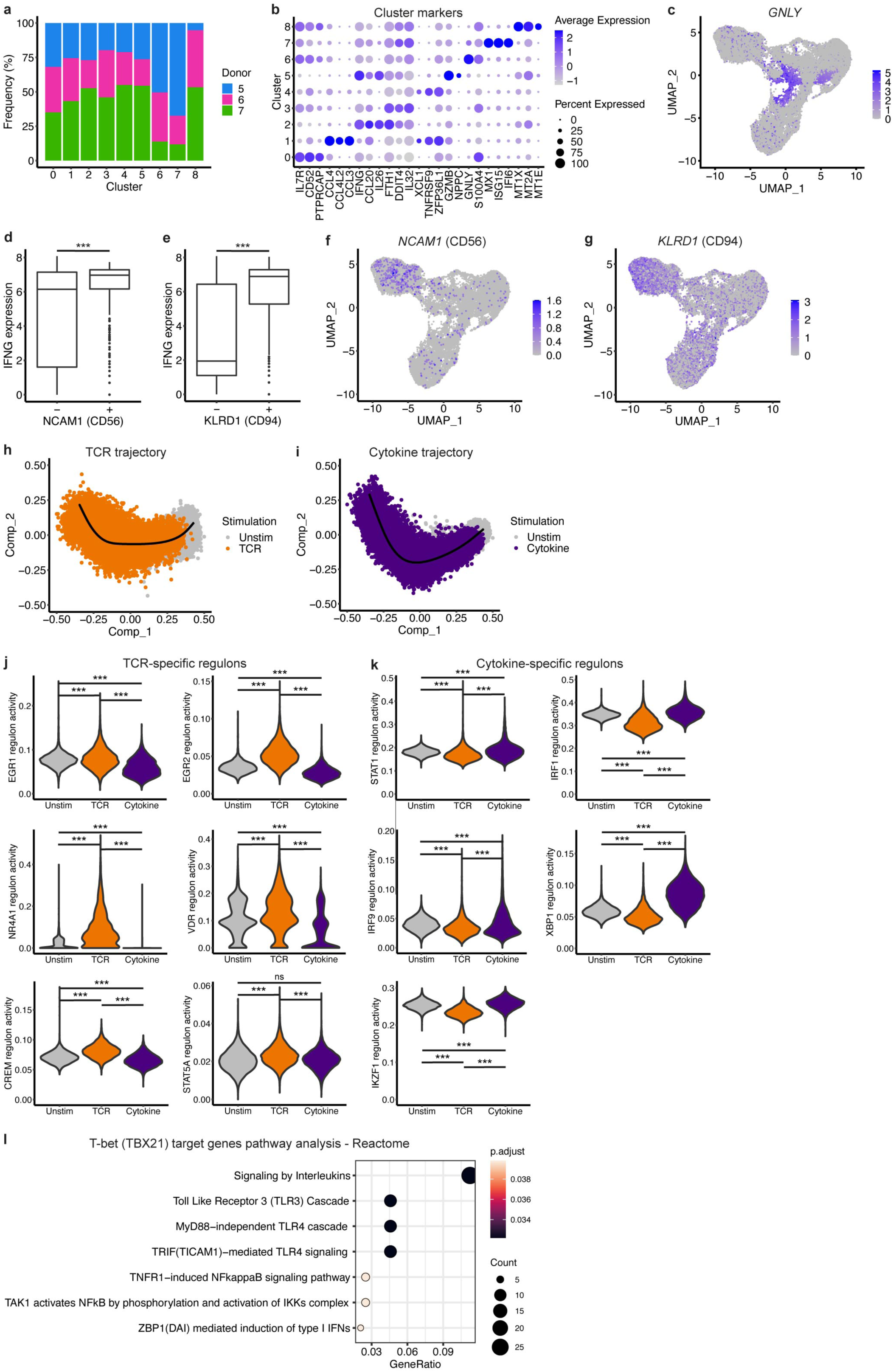
Transcriptional and regulatory analysis of TCR- and cytokine-stimulated MAIT cells. **a**, Proportion of cells from each donor in each cluster (unstimulated, TCR-stimulated and cytokine-stimulated MAIT cells combined). **b**, Expression of the top three marker genes per cluster. Dot size indicates the percentage of cells expressing the gene and dot colour indicates the level of expression. **c**, UMAP coloured by *GNLY* expression. **d**,**e**, Expression of *IFNG* in cytokine-stimulated MAIT cells negative and positive (log-transformed normalised expression > 0) for expression of *NCAM1* (CD56) (**d**) and *KLRD1* (CD94) (**e**). Wilcoxon rank-sum test, *** p < 0.001. **f**,**g**, UMAPs coloured by the expression of *NCAM1* (CD56) (**f**) and *KLRD1* (CD94) (**g**). **h**,**i**, Multidimensional scaling plots of unstimulated and TCR-stimulated (**h**) or unstimulated and cytokine-stimulated (**i**) MAIT cells. SCORPIUS TCR (**h**) and cytokine (**i**) trajectories are shown in black. **j**,**k**, Violin plots showing the activity (AUCell scores) of selected TCR-specific (**j**) and cytokine-specific (**k**) transcription factor regulons in unstimulated, TCR-stimulated and cytokine-stimulated MAIT cells. Differential activity analysis was performed using MAST, *** p < 0.001, ns = non-significant. **l**, Pathway analysis on predicted T-bet (*TBX21*) target genes. Significant Reactome pathways are shown.

**Extended Data Fig. 8.**
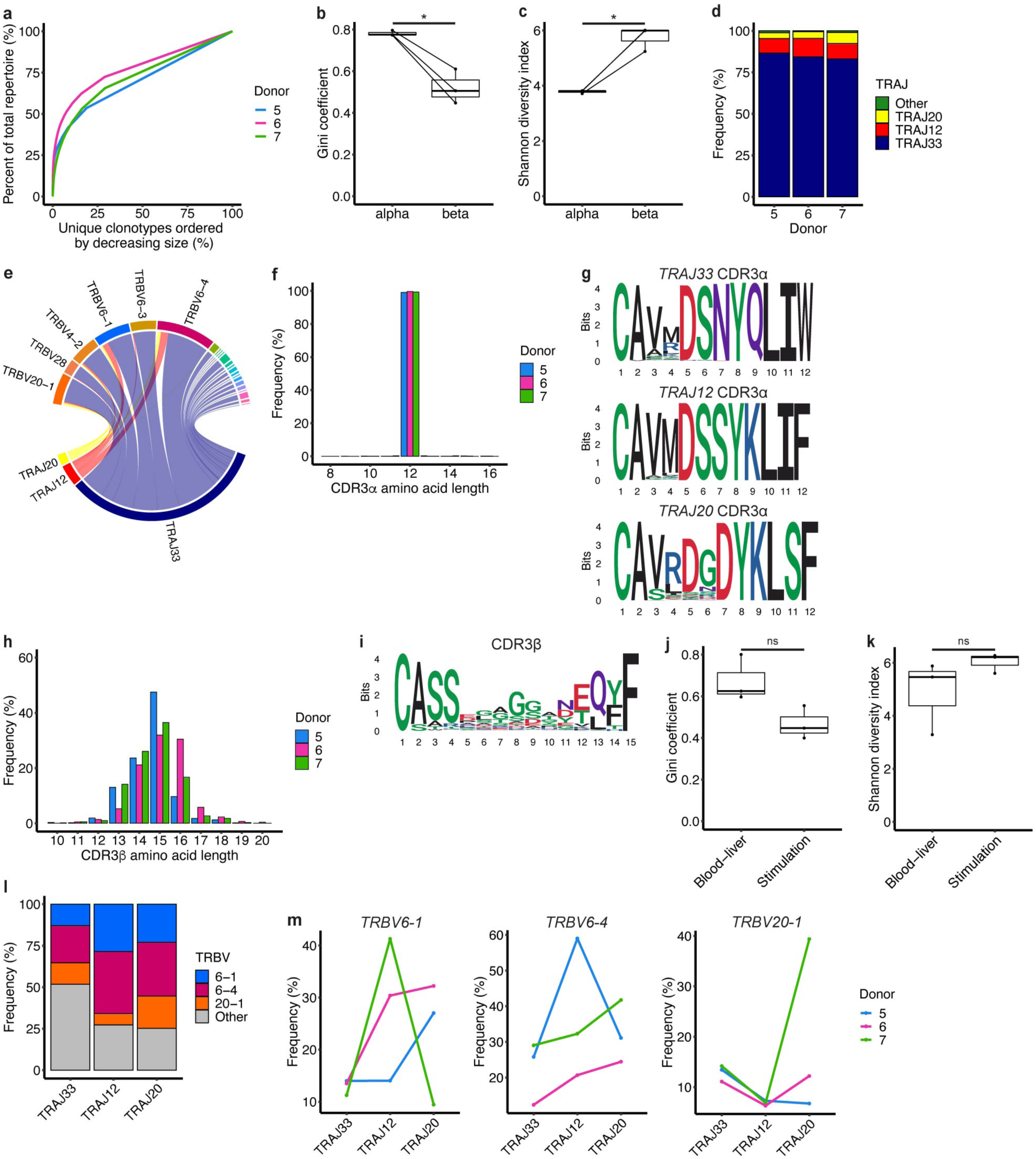
The MAIT cell TCR repertoire in the stimulation experiment is comparable to the blood-liver experiment. TCR repertoire characterisation was performed for unstimulated MAIT cells only. **a**, Line plot demonstrating the clonality of the MAIT cell TCRαβ repertoire in each donor. **b**,**c**, Gini coefficient (**b**) and Shannon diversity index (**c**) for MAIT cell TCRα only and TCRβ only clonotypes. Paired t-test, * p < 0.05. **d**, Proportion of MAIT cells expressing *TRAJ33*, *TRAJ12*, *TRAJ20* and other *TRAJ* gene segments in each donor. **e**, Circos plot showing the average use of *TRAJ* and *TRBV* gene segments amongst MAIT cells combined from all three donors. **f**, MAIT cell CDR3α amino acid length in each donor. **g**, Sequence logos generated from all MAIT cell *TRAJ33*, *TRAJ12* or *TRAJ20* CDR3α amino acid sequences of length 12. **h**, MAIT cell CDR3β amino acid length in each donor. **i**, Sequence logo generated from all MAIT cell CDR3β amino acid sequences of length 15. **j**,**k**, Gini coefficient (**j**) and Shannon diversity index (**k**) for MAIT cell TCRαβ clonotypes in the blood-liver and stimulation experiments. Paired t-test, ns = non-significant. **l**, Proportion of *TRAJ33*, *TRAJ12* and *TRAJ20* MAIT cell TCRs with *TRBV6-1*, *TRBV6-4*, *TRBV20-1* and other *TRBV* gene segments. **m**, Percentage use of *TRBV6-1* (left), *TRBV6-4* (middle) and *TRBV20-1* (right) *TRBV* gene segments amongst *TRAJ33*, *TRAJ12* and *TRAJ20* MAIT cell TCRs. Friedman test with Dunn’s post-hoc test. No significant pairwise comparisons.

**Extended Data Fig. 9.**
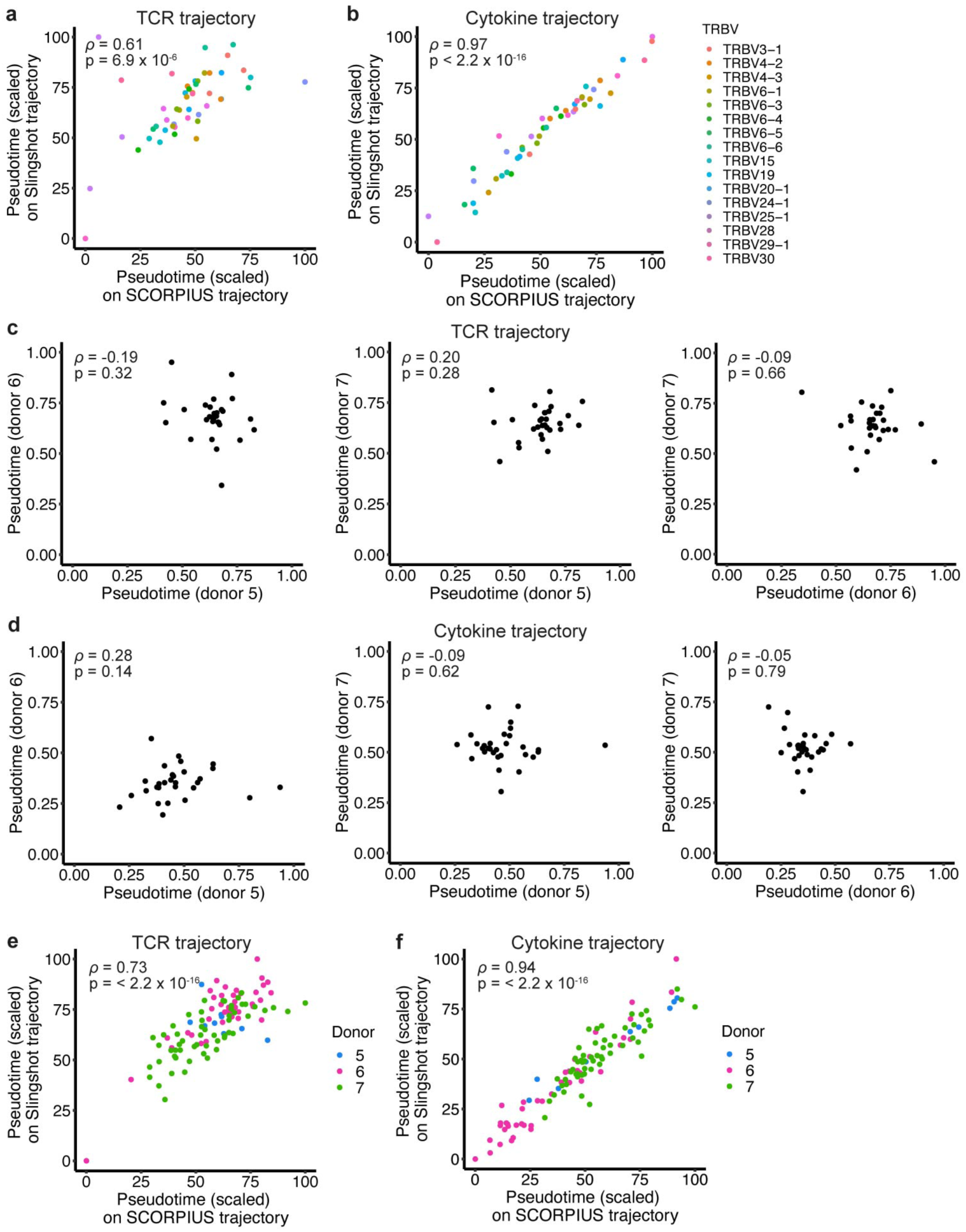
Influence of the TCR repertoire on MAIT cell activation potential. **a**,**b**, Correlation in average *TRBV* pseudotimes on the SCORPIUS and Slingshot (two trajectory methods) TCR (**a**) and cytokine (**b**) trajectories. Values scaled between 0 and 100. **c**,**d**, Correlation in average *TRBV* pseudotimes between pairs of donors on the SCORPIUS TCR (**c**) and cytokine (**d**) trajectories. **e**,**f**, Correlation in average clonotype pseudotimes on the SCORPIUS and Slingshot TCR (**e**) and cytokine (**f**) trajectories. Values scaled between 0 and 100. Plots show stimulated cells only, *TRBV* gene segments with a frequency > 1% in any donor, and clonotypes containing > 20 cells. Spearman’s rank correlation in **a**-**f**.

